# Multifaceted Atlases of the Human Brain in its Infancy

**DOI:** 10.1101/2022.03.19.484985

**Authors:** Sahar Ahmad, Ye Wu, Zhengwang Wu, Kim-Han Thung, Weili Lin, Gang Li, Li Wang, Pew-Thian Yap

## Abstract

Brain atlases agglomerate structural and functional features of a population of individuals in a standard coordinate space. Here, we introduce for the first time a collection of atlases that charts postnatal development of the human brain in a spatio-temporally dense manner from two weeks to two years of age. Atlases capturing month-to-month changes of the human brain are essentially nonexistent for the first two years of life — the critical developmental period during which the brain is evolving in virtually all facets at an exponential pace. This unmet need is compounded by the lack of atlases that provide a unified and holistic picture of the brain from both the perspectives of cortical surface convolutions and tissue volumetric characteristics. Existing surface and volumetric atlases are typically constructed independently in different coordinate spaces, causing discrepancies and complications in multifaceted analyses. Our month-specific conjoint surface and volumetric atlases chart normative patterns and capture key traits of early brain development and are therefore critical for identifying aberrations from normal developmental trajectories. Our atlases represent a major advance toward providing the neuroscience community an invaluable resource to facilitate the understanding of early structural and functional development by mapping multiple measurements of infant brains in a common reference frame for precise spatio-temporal quantification of cortical and subcortical changes.

## Introduction

**H**uman brain development is a complex, protracted process that begins during the third gestational week and extends through adulthood^1^. Throughout late prenatal period and early infancy, the human brain undergoes a myriad of cellular processes, including neurogenesis, neuronal migration, astrogliogenesis, oligodendrogenesis, synaptogenesis, myelination, and synaptic pruning^2^. These cellular events drive the remarkably rapid growth of the infant brain during the first two years of life, resulting in drastic structural changes and reorganization of neural circuits^3–5^. The intracranial volume of the newborn’s brain doubles during the first postnatal year, attaining approximately 65% of the adult brain volume^6^. The gray matter (GM) volume increases more dramatically (∼71%) than the white matter (WM) volume (∼20%) in the first year. The cerebellum grows at the fastest rate and more than doubles in volume during the first 90 days after birth. In contrast, the hippocampus grows at the slowest rate by only 47% during the same period^7^. A more precise quantification of early brain growth will be conducive to a step change in our understanding of developments that lead to maturation of cognitive functions^3^. However, endeavors in this direction have so far been hindered by the dearth of brain atlases for mapping features of early brain development to common spaces necessary for fine-grained spatio-temporal quantification of brain changes.

Unlike the long-established growth charts utilized by pediatricians to quantify year-to-year maturation in terms of a child’s height, weight, and head circumference in relation to standardized curves derived from healthy growing children^8, 9^, normative references for neurodevelopment are virtually nonexistent. Recent efforts have been dedicated toward constructing brain charts that capture normative patterns of cerebral development^6^ in terms of volumetric growth of GM, WM, and cerebrospinal fluid (CSF), and cortical growth captured by surface area and cortical thickness^10^. Although these brain charts standardize brain morphological measurements, they do not define a common coordinate system for mapping brain features, particularly from multiple imaging modalities, to a unified reference space to form a more complete picture of brain development. An additional limitation of existing brain charts is that they rely on the demarcation of the brain into adjoint but separate brain areas with hypothetically uniform functions, while in reality an extensive body of evidence suggests a more gradual transition of areal boundaries, particularly in the association cortices^11–13^. These limitations call for the construction of surface-volume atlases that offer a standardized coordinate framework for concurrent analysis of multimodal data with fine granularity at the voxel or vertex level.

Existing atlases of the baby brain are limited to specific prenatal or postnatal periods^14–26^ (Supplementary Fig. 1). Brain atlases densely covering the entirety of the first two years of life are sorely lacking owing to the unique challenges in collecting longitudinal brain MRI data. Adding to the difficulties in constructing these much needed atlases are the rapid changes in the sizes and shapes of cerebral structures (Supplementary Fig. 2), and the evolving and often insufficient tissue contrast between WM and GM^27^. Moreover, existing cortical surface and volumetric atlases are typically constructed independently, resulting in the following problems: (i) misalignment between tissue interfaces of the volumetric atlas and the white and pial surfaces of the cortical atlas. (ii) Fuzzy cortical structures, as volumetric displacements are estimated without taking into account the complex convolutions of the cortical surface. (iii) Anatomically implausible displacements owing to cortical surface registration based only on surface attributes but ignoring the volumetric data^28^. (iv) Volumetric and cortical surface attributes are not defined in the same space, making it difficult to analyze the two entities concurrently and consistently^29^. (v) Inconsistent and inaccurate alignment may diminish real but subtle changes^30^.

In this work, we present for the first time a set of month-specific surface-volume longitudinal brain atlases of infants from 2 weeks to 24 months of age. These atlases facilitate the precise mapping of fine-grained changes in both space and time, and are therefore a valuable resource for studying postnatal human brain development, identifying early roots of neurological disorders, and quantifying development-related malformations. We jointly construct the surface and volume atlases for consistent alignment of WM-GM and GM-CSF interfaces in both surface and volumetric spaces, making possible unified surface and volumetric analyses in a single coordinate space, facilitating the investigation of the interplay of developmental trajectories of multiple brain features. We demonstrate that our high-fidelity atlases capture the normative evolutionary landscape of cortical surface features and tissue volumetric characteristics of the infant brain during the first two years of critical brain development.

## Results

### Surface-volume consistency

Our atlases allow multifaceted analyses of volumetric and surface data in a common space for holistic and unified understanding of early brain development. As a reference, we first construct the volumetric and surface infant brain atlases at 12 months, IBA12-V and IBA12-S, in a surface-volume consistent manner^31^ using high-quality magnetic resonance (MR) images of 37 infant subjects scanned around 12 months of age as part of the Baby Connectome Project (BCP)^32^. The cortical surface and volumetric data of these subjects are simultaneously normalized in space^33^ and then fused to form (i) the white and pial cortical surfaces of IBA12-S and T1-weighted (T1w) intensity image, T2-weighted (T2w) intensity image, and (ii) the WM, GM, and CSF tissue maps of IBA12-V for rich characterization of brain morphology. We will show that IBA12-V and IBA12-S yield substantial advantages over the cortical surface atlas constructed with Spherical Demons^34^ and the volume and surface atlases constructed with ANTs^35^.

In contrast to atlases constructed with Spherical Demons and ANTs, IBA12-V and IBA12-S exhibit remarkably sharp anatomical details despite being an average of a few tens of subjects (Fig. 1), thanks to the good alignment of GM-WM boundaries via explicit registration of cortical surfaces. In fact, IBA12-V resembles MR images of one-year-olds with substantially less fuzziness than the atlas constructed with ANTs without leveraging the geometry of cortical surfaces. This is evident from the close-ups of the temporal and parietal lobes (Figs. 1a,b). Subcortical structures, including the thalamus, caudate nucleus, and putamen, are well defined, indicating that the surface-volume consistent atlas construction process is conducive to preserving both cortical and subcortical anatomical details^36^. This is not surprising because the putamen, for example, are only separated from insular cortex by a narrow band of WM. While the T1w IBA12-V exhibits adult-like contrast, the T2w IBA12-V is close to isointense with significant overlap in intensity distributions between WM and GM, consistent with T2w images of one-year-olds^37^. This is particularly noticeable at the superior frontal and rostral middle frontal cortices (Fig. 1b).

**Fig. 1.**
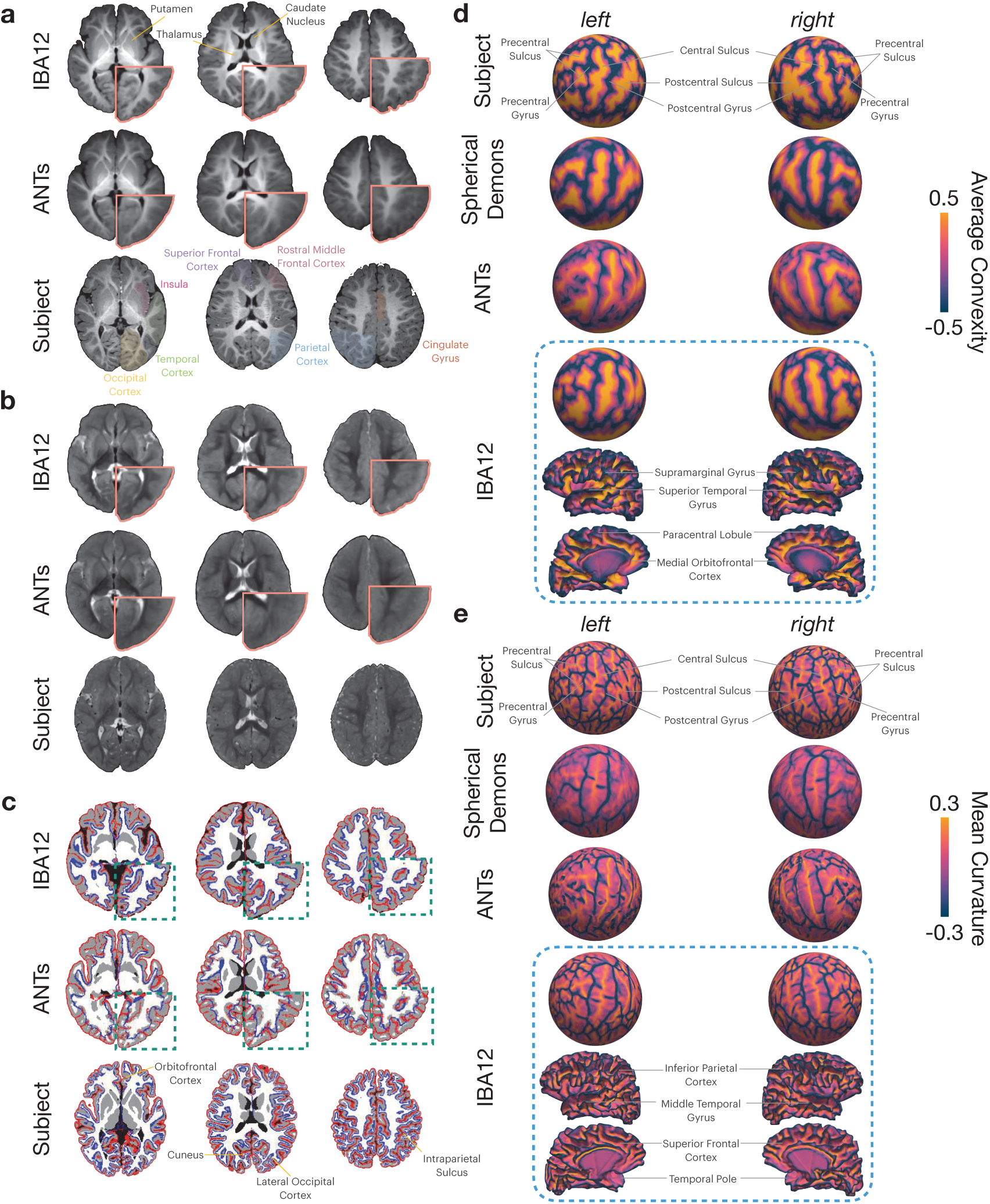
The reference infant brain atlas at 12 months. **a,b**, Transverse sections of the T1w and T2w atlases, and the individual subject scan. **c**, White and pial cortical surface atlases overlaid onto the tissue segmentation maps of the atlases. The cortical surfaces of individual subject are also projected onto its tissue segmentation map. **d**, Average convexity mapped onto the spherical surfaces of the subject and the surface atlases. Lateral and medial views of the white surface of IBA12-S color-coded by average convexity are also shown. **e**, Mean curvature mapped onto the spherical surfaces of the subject and the surface atlases. Lateral and medial views of the white surface of IBA12-S color-coded by mean curvature are also shown.

The white and pial cortical surfaces of IBA12-S are consistently aligned with the GM-WM and WM-CSF interfaces as delineated by the tissue segmentation maps of IBA12-V (Fig. 1c). This is confirmed by the zoomed-in views of, for example, the lateral occipital, inferior parietal, cuneus, precuneus, medial orbitofrontal, and superior parietal cortices. On the other hand, the ANTs atlas shows inconsistent alignment of the cortical surfaces with tissue interfaces in the volumetric space.

The cortical surfaces of both hemispheres in IBA12-S preserve cortical convolutions with distinct gyral and sulcal patterns (Figs. 1d,e). For greater clarity, we map the average convexity and mean curvature of the white surface onto a sphere^1^. Average convexity captures coarse-scale geometric features of primary sulcal folds^38^, whereas mean curvature captures fine-scale geometric features of secondary and tertiary folds. The average convexity maps of the Spherical Demons and IBA12-S atlases are consistent with that of an individual subject. On the other hand, the average convexity map of the ANTs atlas indicates atypical narrowing of the gyral and sulcal folds. Only the IBA12-S exhibits typical mean curvature patterns. Spherical Demons gives a fuzzy mean curvature map that fails to capture fine-grained details of cortical folds. ANTs gives an atypical mean curvature map with holes, indicating alteration of surface topology. These results indicate that primary gyri and sulci, including the precentral and postcentral gyri and sulci and the central sulcus, are captured by all three methods, but localized gyral and sulcal details are only preserved in the IBA12-S.

### Surface atlases across infancy

Early brain development is characterized by dynamic changes in cortical folding patterns. We constructed month-specific cortical atlases for the first two years of life by longitudinally warping the IBA12-S. As shown in Fig. 2a, major cortical folds of white surfaces — including the central sulcus, precentral and postcentral gyri, inferior parietal lobule, temporal gyrus, superior and inferior temporal sulci, superior frontal gyrus, cingulate gyrus and sulcus, and parieto-occipital sulcus — are highly consistent across time points. Mean curvature mapped onto the inflated white surface atlases indicates subtle developments of secondary and tertiary gyri and sulci (Fig. 2b). Cortical thickness measured between the white and pial cortical surfaces of the atlases, projected onto the inflated white surface atlases (Fig. 2c), indicates that the cortical thickness of the prefrontal cortex, temporal lobe, and insula increases dramatically. In contrast, the thickness of the visual and sensorimotor cortices increases at a slower pace. Month-to-month surface atlases are provided in Supplementary Figs. 3-5.

**Fig. 2.**
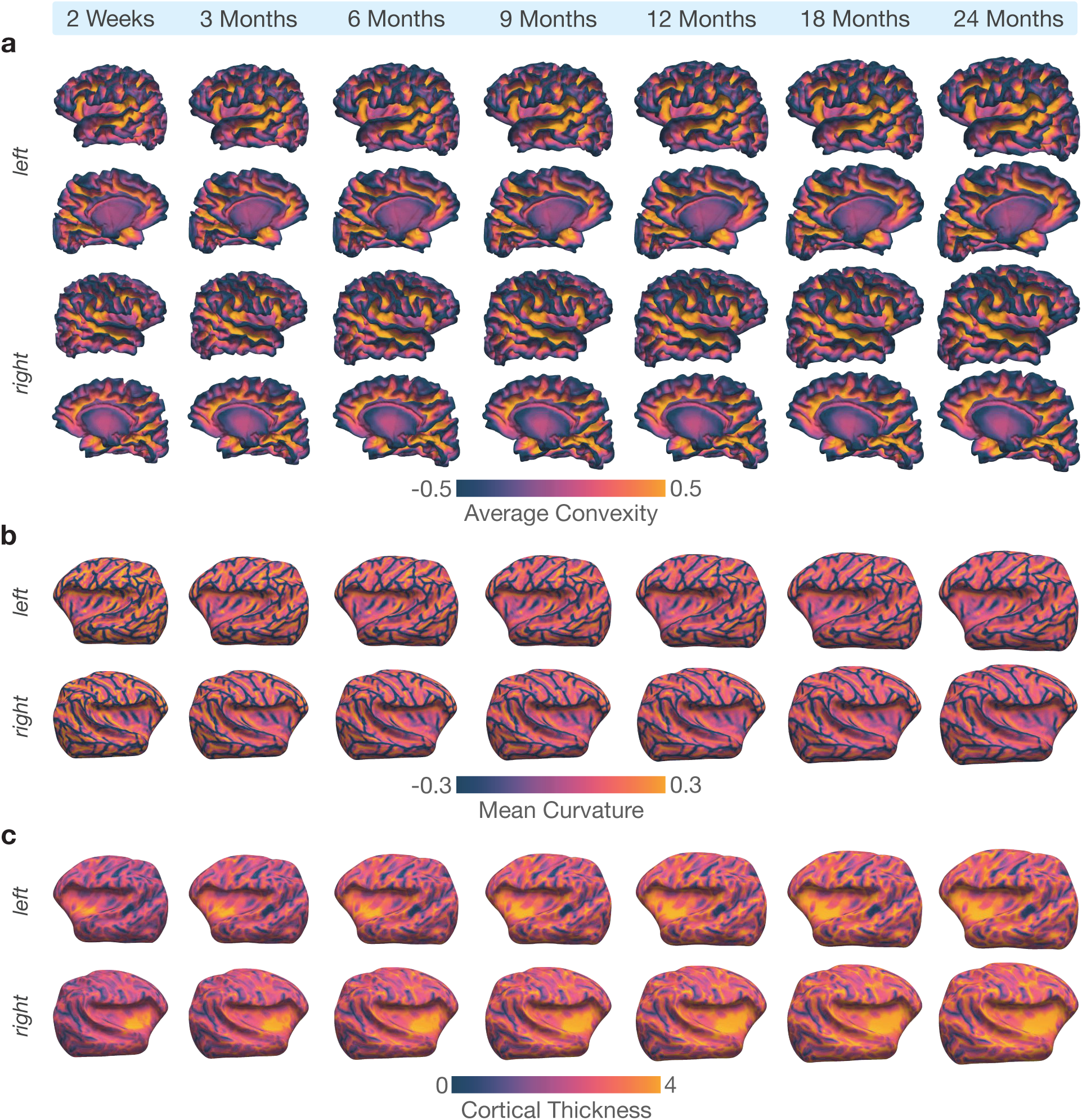
Longitudinal white surface atlases of the infant brain. **a**, Lateral and medial views of the cortical surface atlases of the white surfaces from 2 weeks to 24 months. The cortical surface atlases are color-coded by their respective average convexity (millimeter) maps. **b**, Mean curvature (millimeter^−1^) map projected onto the inflated white surface atlases at selected time points from 2 weeks to 24 months. **c**, Cortical thickness (millimeter) mapped onto the inflated white surface atlases at selected time points from 2 weeks to 24 months.

### Volumetric atlases across infancy

The IBA-V, T1w and T2w atlases, from 2 weeks to 24 months are presented in Figs. 3a,b and Supplementary Figs. 6 and 7. The image contrast evolves month-to-month during year one and becomes adult-like in year two. The T1w and T2w atlases are isointense around 3 – 6 months and 9 – 12 months, respectively. The atlases preserve distinct boundaries at the GM-WM interface as evident from the close-ups of the precuneus, cuneus, and inferior parietal cortex (Figs. 3a,b).

**Fig. 3.**
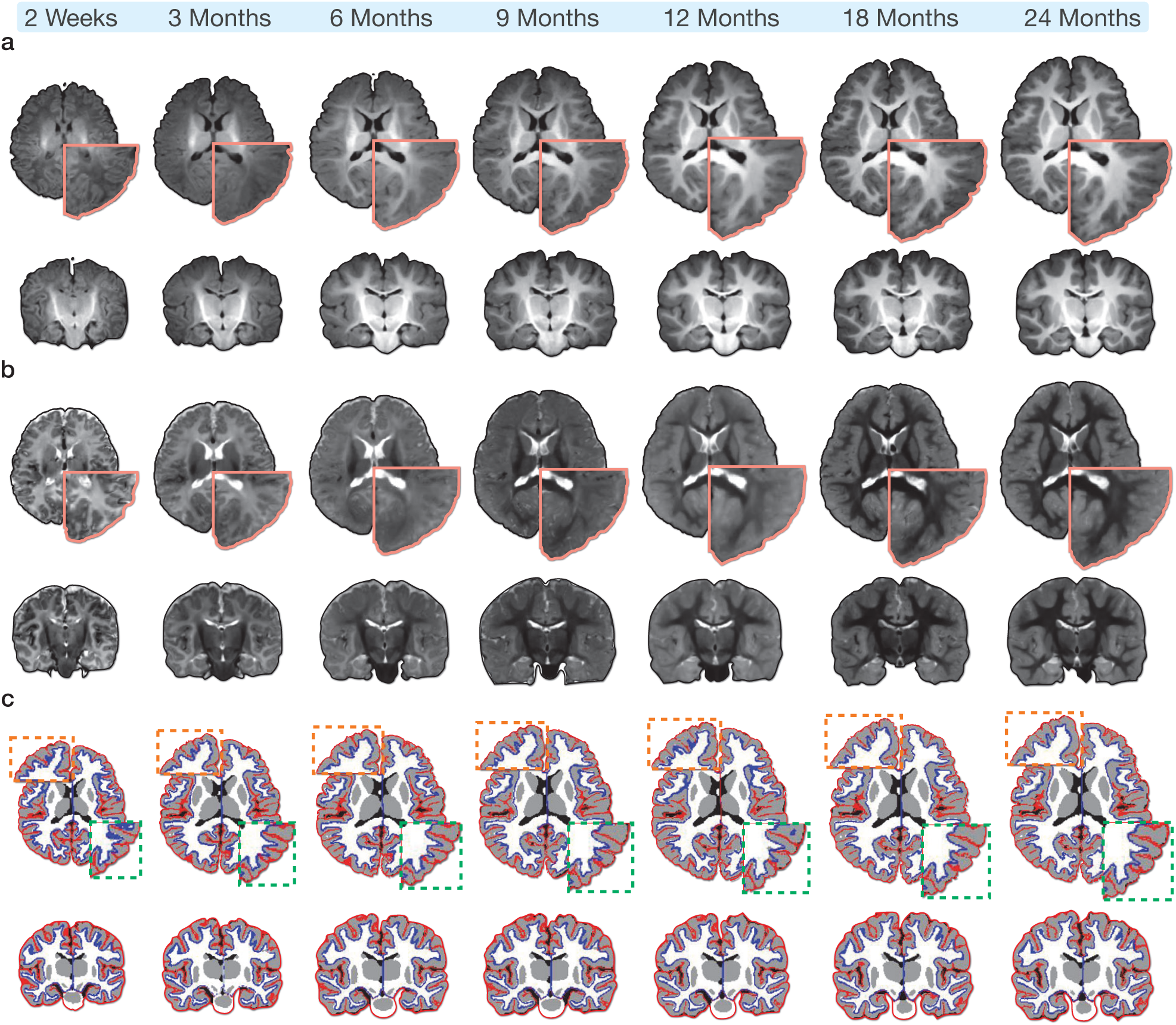
Volumetric atlases of the infant brain. **a,b**, Transverse and coronal sections of the T1w and T2w atlases from 2 weeks to 24 months, respectively. **c**, Cortical atlases of the white (*blue*) and pial (*red*) surfaces overlaid onto the corresponding volumetric atlases at selected time points between 2 weeks and 24 months. The close-ups of the frontal and parietal lobes show that the cortical surface atlases are well-aligned at the WM-GM and GM-CSF interfaces.

Age-specific white and pial cortical surface atlases overlapped on the tissue map atlases (Fig. 3c and Supplementary Fig. 8) indicate that the cortical surface atlases are well aligned with the tissue boundaries of the volumetric atlases, particularly at the superior and middle frontal gyri of both the hemispheres, supramarginal gyrus, inferior parietal cortex, insula, and precentral and postcentral gyri. Good alignment is confirmed by zoomed-in views of the atlases at selected time points (Fig. 3c).

### Surface and volumetric development

The IBA captures the developmental features of typically-developing infants in the first two years of life. Compared with the atlases constructed with Spherical Demons and ANTs, the IBA more faithfully reflects the individuals, as evidenced by the smaller root mean square error (RMSE) between the growth curve for each surface or volumetric feature of the IBA and the population curve obtained by fitting a generalized additive mixture model (GAMM)^39^ (Fig. 4a).

**Fig. 4.**
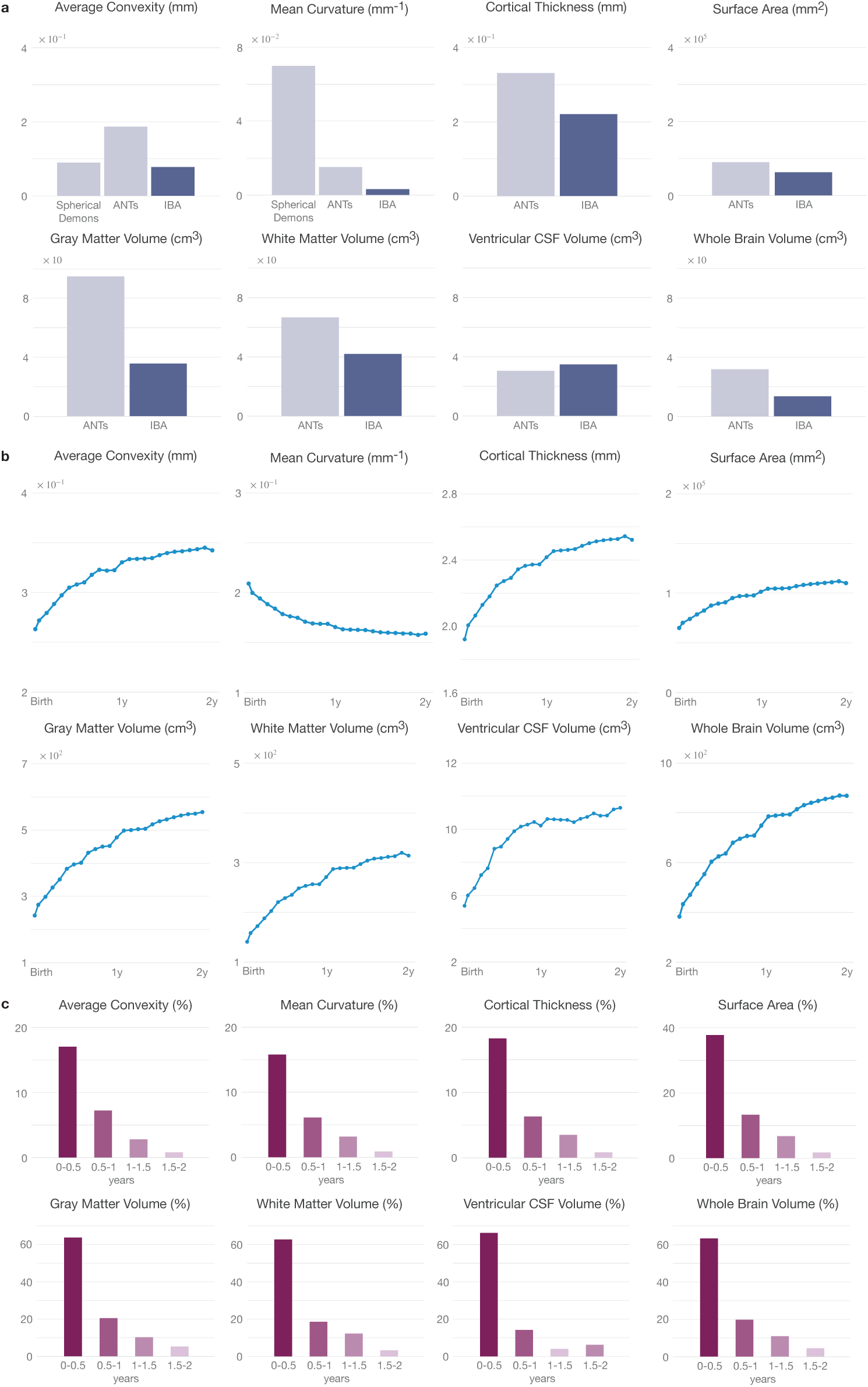
Development of surface and volume. **a**, RMSE of the developmental trajectories of surface and volumetric features, computed between the atlases and the individual subjects. **b**, Developmental trajectories of white surface features and tissue volumes of the IBA. **c**, Surface and volumetric changes every six months.

We studied the surface and volumetric features captured by the IBA (Figs. 4b,c). During the first and second postnatal years, the IBA-S increases^2^ in average convexity by 25.5% and 3.6%, decreases in mean curvature by 20.9% and 3.9%, increases in cortical thickness by 25.7% and 4.3%, and increases in surface area by 56.1% and 8.5%. During the same period of time, the IBA-V increases in GM volume by 97.2% and 13.8%, increases in WM volume by 92.9% and 15.9%, increases in ventricular CSF volume by 89.8% and 10.5%, and increases in whole brain volume (WBV = GM + WM) by 95.6% and 14.6%. The velocity curves for these surface and volumetric features are shown in Supplementary Fig. 9.

Regional analysis indicates spatially heterogeneous development. We measured the average convexity of 34 cortical regions-of-interest (ROIs) delineated via FreeSurfer using the Desikan-Killiany atlas^40^. All regions^3^ (Fig. 5) show increasing trend but at varying rates ranging from 19% to 34% and 1.2% to 5.7% for the first and second postnatal years (Fig. 7a), respectively. Average convexity exhibits higher growth rate in association cortices compared with primary cortices. The growth rates (Year 1, Year 2) are (24.2%, 1.5%) for the primary visual cortex, (23.3%, 3.3%) for the primary somatosensory cortex, (22.9%, 3.8%) for the primary motor cortex, (27.0%, 3.6%) for the primary auditory cortex, (27.1%, 4.2%) for the temporal association cortex, (27.1%, 3.6%) for the parietal association cortex, and (23.9%, 3.7%) for the prefrontal association cortex.

**Fig. 5.**
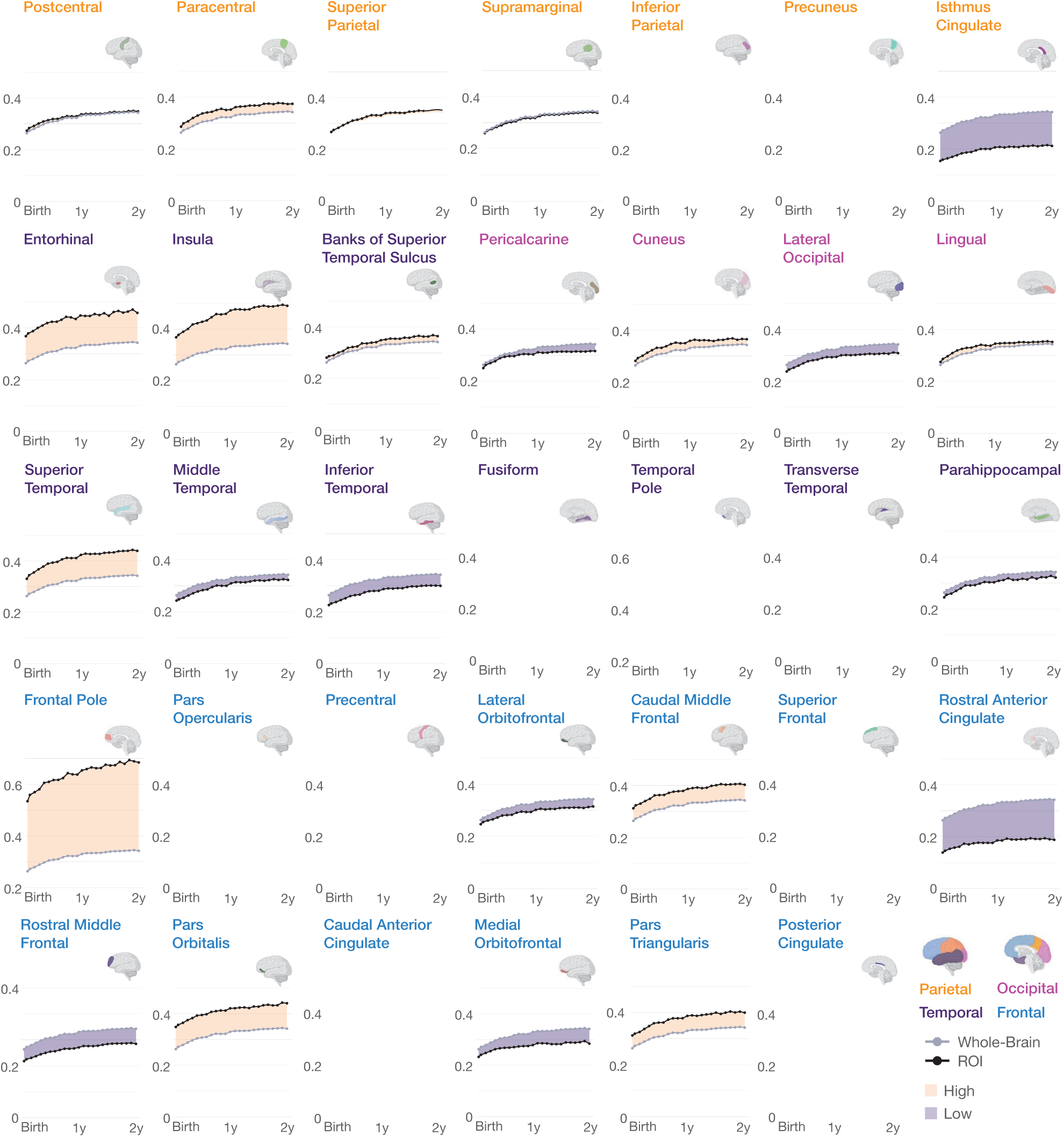
Regional developmental trajectories of average convexity. Growth curves of average convexity for IBA cortical regions. Shaded regions indicate whether average convexity is greater or lower than average.

In terms of mean curvature, all cortical ROIs exhibit decreasing trends (Supplementary Figs. 11, 13a) at varying negative growth rates ranging from 14% to 26% and 1% to 6% in the first and second postnatal years, respectively. The negative growth rates (Year 1, Year 2) are (19.9%, 2.0%) for the primary visual cortex, (19.5%, 3.9%) for the primary motor cortex, (21.0%, 3.4%) for the primary somatosensory cortex, (20.6%, 3.7%) for the primary auditory cortex, (21.5%, 3.6%) for the temporal association cortex, (21.2%, 4.1%) for the parietal association cortex, and (18.5%, 4.1%) for the prefrontal association cortex.

Developmental trajectories of regional cortical thickness in Supplementary Fig. 14 show that the insula, frontal, superior, middle, and inferior temporal cortices are consistently thicker during infancy. On the other hand, the precentral and postcentral gyri, occipital and inferior parietal cortices, and banks of superior temporal suclus are thinner throughout the study period. Regional growth rates vary from 21.2% to 32.0% and 2.3% to 6.7% for the first and second postnatal years (Supplementary Fig. 16a). The growth rates (Year 1, Year 2) are (22.6%, 2.3%) for the primary visual cortex, (25.1%, 3.3%) for the primary somatosensory cortex, (24.5%, 4.3%) for the primary motor cortex, (29.5%, 4.0%) for the primary auditory cortex, (27.7%, 4.4%) for the temporal association cortex, (27.6%, 4.4%) for the parietal association cortex, and (23.9%, 4.8%) for the prefrontal association cortex.

We show the developmental trajectories of regional surface area in Supplementary Fig. 17 and growth rates for the first and second postnatal years in Supplementary Fig. 19a. High-expansion regions include rostral anterior cingulate gyrus, medial orbitofrontal gyrus, superior frontal gyrus, rostral middle frontal gyrus, inferior and middle temporal gyri, and superior and inferior parietal gyri. Low-expansion regions include insula, pericalcarine, cuneus, entorhinal, temporal pole, and parahippocampal gyrus. Regional growth rates vary from 20.5% to 49.7% in year 1 and 2.9% to 11.1% in year 2. The growth rates (Year 1, Year 2) are (42.4%, 3.4%) for the primary visual cortex, (36.0%, 5.6%) for the primary somatosensory cortex, (34.1%, 6.0%) for the primary motor cortex, (42.3%, 5.3%) for the primary auditory cortex, (39.8%, 6.9%) for the temporal association cortex, (38.9%, 5.8%) for the parietal association cortex, and (38.1%, 7.7%) for the prefrontal association cortex.

The IBA captures the asymmetry of cortical features. We show the hemispheric lateralization of regional cortical features via the laterality index, LI = (left − right)/(left + right), computed for each ROI. There is significant asymmetry (*t*-test; *p <* 0.01) in average convexity (Fig. 6), mean curvature (Supplementary Fig. 12), cortical thickness (Supplementary Fig. 15), and surface area (Supplementary Fig. 18) for all ROIs. The corresponding *t*-scores and degrees of freedom (DoFs) are reported in Supplementary Tables 1, 2, 3, 4. The ROI-specific mean LI for average convexity, mean curvature, cortical thickness, and surface area are shown in Fig. 7b and Supplementary Figs. 13b, 16b, 19b.

**Fig. 6.**
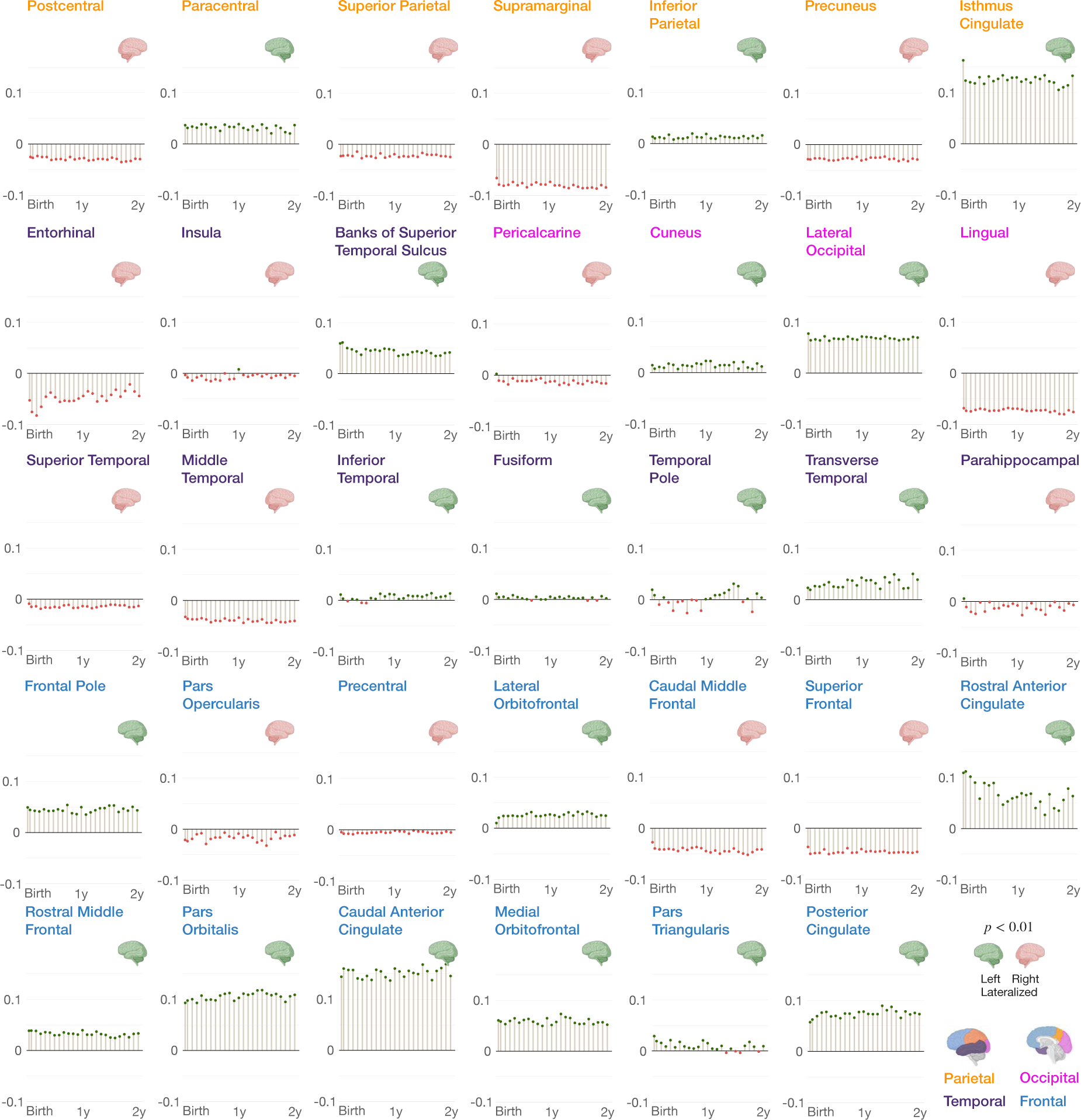
Hemispheric asymmetry of average convexity. Region-specific laterality index for average convexity of the IBA. Positive laterality is associated with left lateralization (*p <* 0.01) and negative laterality is associated with right lateralization (*p <* 0.01).

**Fig. 7.**
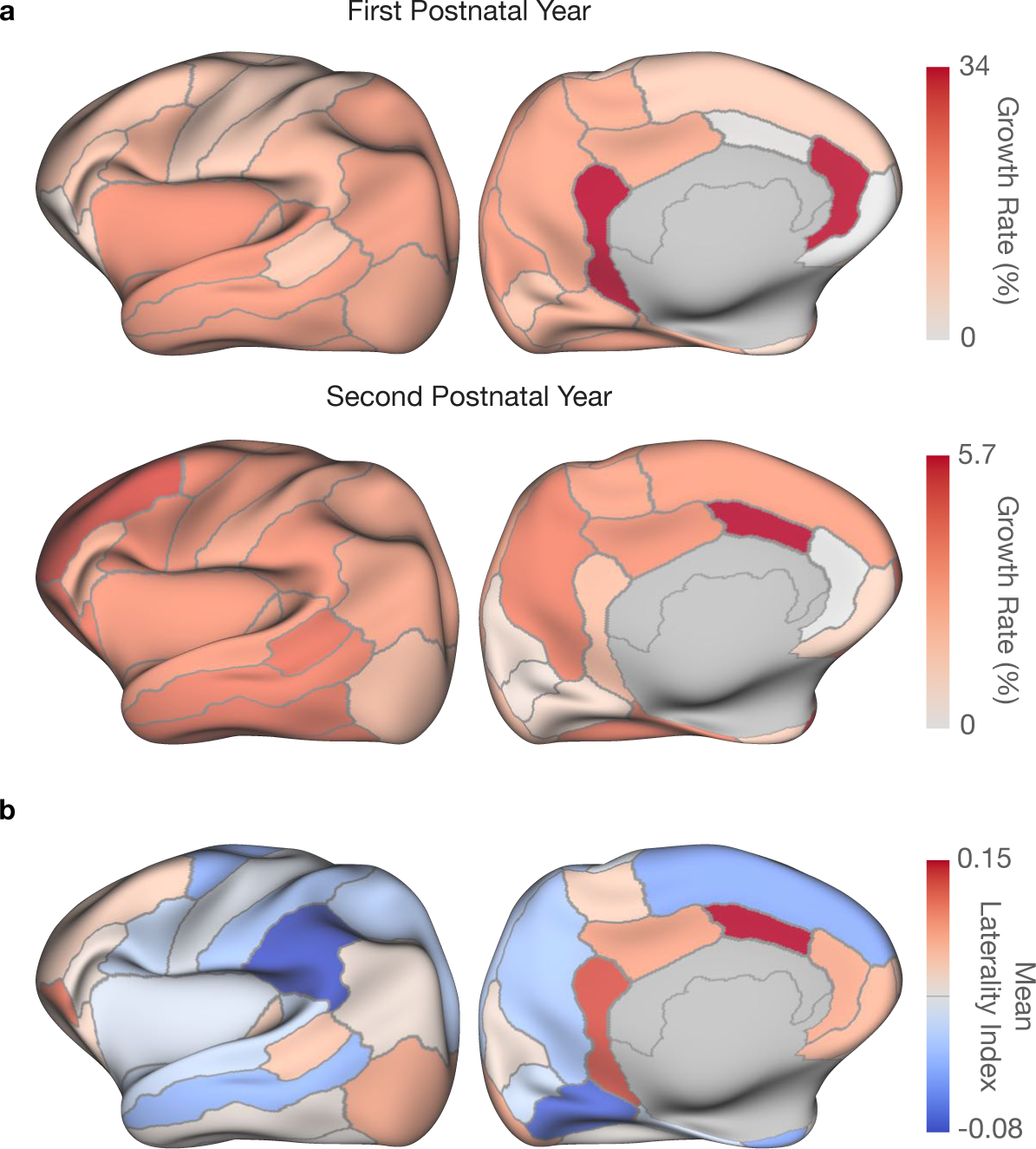
Analysis of average convexity. **a**, Regional growth rates in terms of average convexity for the first (*top row*) and second (*bottom row*) postnatal years. **b**, ROI-specific mean laterality index for average convexity.

### Cortical myelination

We investigated cortical myelination across infancy by T1w/T2w ratio mapping of the cortical ribbon to the white surfaces of individual subjects^41^. The GAMM-fitted myelin maps (Fig. 8) indicate increasing myelination in both cerebral hemispheres at average rates of 57.7% and 7.9% in the first and second postnatal years, respectively. Myelination is spatially heterogeneous with heavy myelination in the primary viusal, motor, and somatosensory cortices and light myelination in the prefrontal, parietal, and temporal association cortices.

**Fig. 8.**
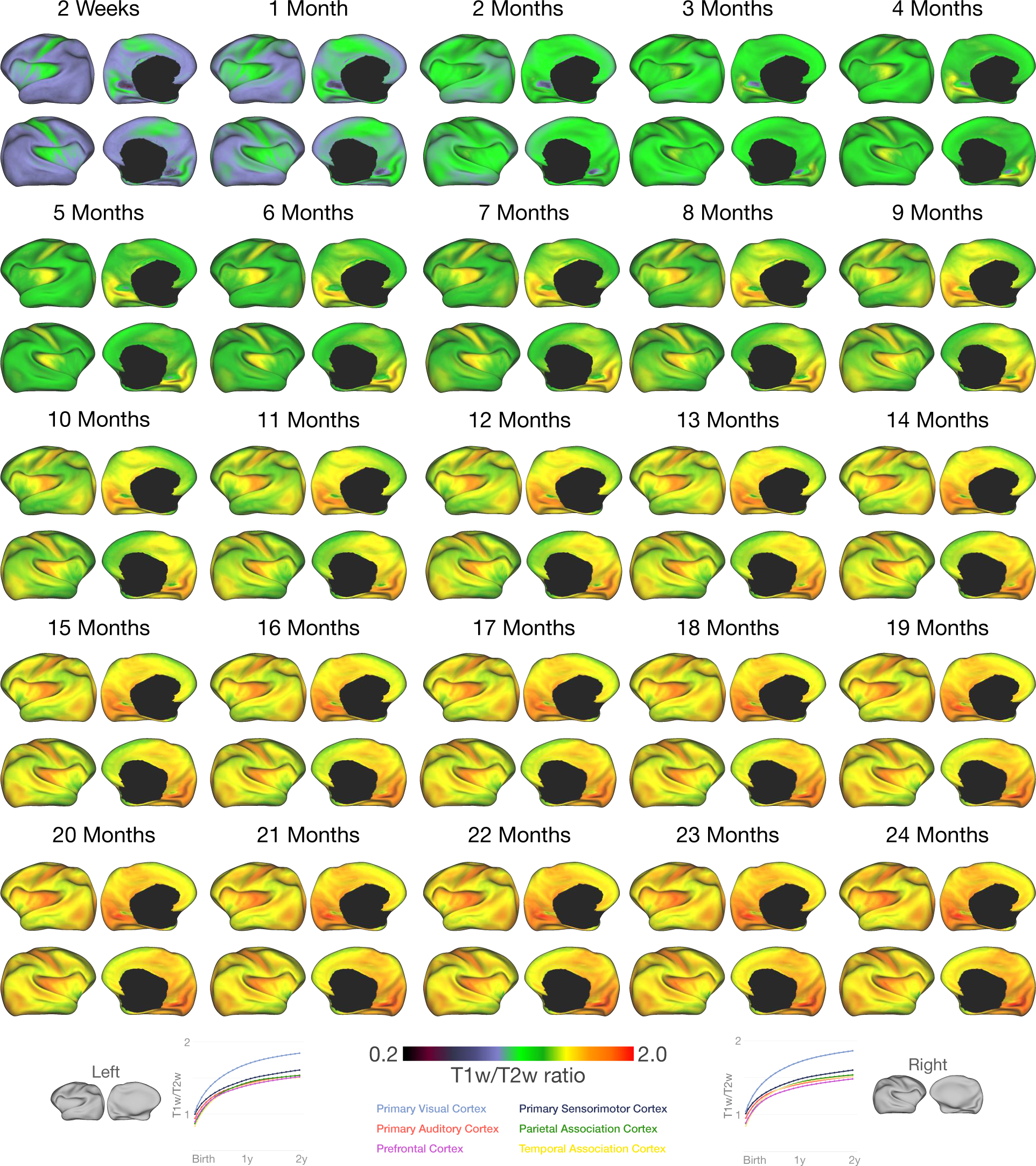
Cortical myelination across infancy. Cortical T1w/T2w ratio mapped onto the inflated white surfaces of the IBA. Myelination is heterogeneous across cortical regions.

## Discussion

Charting normative structural and functional changes during early brain development is key to understanding aberrations associated with neurodevelopmental disorders such as attention-deficit/hyperactivity disorder (ADHD), dyslexia, and cerebral palsy^42–44^. To this end, atlases provide a basis for precise comparison of normative and aberrant development. We have presented for the first time a set of longitudinal surface-volume atlases of the infant brain covering every month of the first two postnatal years. These atlases represent an unprecedented resource that will facilitate the joint investigation of changes in cortical geometries and brain tissues during a period of rapid and critical brain development.

Atlases are commonly used to spatially normalize MRI data for quantification of anatomical volumes and investigation of structural/functional connectivity and cortical macrostructure/ microstructure^45–49^. However, existing infant atlases (Supplementary Table 5 and Supplementary Fig. 1) are constructed independently either in the surface space or the volumetric space, causing inconsistencies between cortical surfaces and tissue boundaries. We resolved this issue by conjointly constructing cortical surface and volumetric atlases in a common space. Our atlases consist of T1w and T2w intensity images, tissue segmentation maps of GM, WM, and CSF, and white and pial surfaces for every month between 2 weeks and 24 months. Our atlases are substantially better in preserving anatomical details and surface geometric features than ANTs ^35, 50, 51^ and Spherical Demons^34, 52^.

The human brain undergoes complex gyrification that begins after mid-gestation^53, 54^, nudging the initially smooth cerebral surface into a highly convoluted structure. Several hypotheses have been put forward to explain the mechanics driving gyrification, including axonal tension^55^, tangential expansion of the cortical layers relative to sublayers generating stress^56^, and biochemical prepatterning of the cortex^57^. Although the sizes and shapes of gyri and sulci vary across individuals, primary gyri and sulci share common orientations and locations^58^. Primary folds are associated with motor and sensory cortices, which mature relatively early even at birth and continue to refine postnatally. The IBA faithfully retains major gyral and sulcal folds.

Average convexity quantifies the degree of folding of primary gyri and sulci. Region-specific growth trajectories of average convexity yielded by the IBA reveal that the temporal association cortex evolves at the fastest rate, followed by the parietal and frontal cortices. Primary sensorimotor and visual cortices grow at a slower rate. This suggests that the primary cortices are already developed at term birth; therefore, they tend to grow slower compared with association cortices, which are involved in high-order functions and are immature at birth and undergo protracted development^59^.

The IBA shows inter-hemispheric differences in regional average convexity. Left-lateralized regions are mainly located in the frontal and temporal cortices, whereas right-lateralized regions are prominent in the parietal cortex. This suggests that anatomical asymmetry may correspond to functional asymmetry. For instance, the triangular and orbital parts of the inferior frontal gyrus are language-related^60, 61^ and therefore exhibit leftward asymmetry. Similarly, the transverse temporal gyrus is left-lateralized as it is part of the auditory processing network^62^. The frontal pole is responsible for action selection and involves more of the left hemisphere than the right^63^. The lateral occipital cortex and fusiform gyrus, involved in face processing, are left-lateralized^64^. Furthermore, our results confirm that the parietal cortex, comprising supramarginal gyrus, superior parietal cortex, and precuneus, shows right laterality, consistent with the functional dominance of the right hemisphere in visuospatial processing tasks^65^. The postcentral gyrus, which is a part of the somatosensory cortex, exhibits leftward asymmetry, whereas the precentral gyrus, which is involved in voluntary motor movements, shows rightward asymmetry^65^.

The human brain buckles and folds at different extents^66^. At a larger scale, it curves into a spheroid split into two hemispheres. At a smaller scale, it carves into primary, secondary, and tertiary gyri and sulci in association with cortical development. The degree of small-scale cortical folding is quantified via mean curvature that measures the extrinsic curvature of the cortical surface^67^. The IBA shows that the absolute mean curvature decreases substantially during the first year and stabilizes in the second year. The decrease in mean curvature is due to the dynamic cortical folding associated with post-migrational processes characterized by axogenesis, dendritogenesis, synapse initiation, maturation, and pruning of both pyramidal neurons and cortical interneurons^37^. Morphologically, the downward trend in mean curvature reflects the widening of sulci due to the increase in brain size, resulting in sulcal fundi and gyral crowns that curve less sharply. Mean curvature exhibits differential decrease in all cortical ROIs: entorhinal cortex and temporal pole show fastest decline, whereas rostral and caudal anterior cingulate cortices show slowest decrease. Asymmetry of mean curvature is significant in all regions. In line with Remer et al.^68^, the posterior cingulate and rostral anterior cingulate cortices are left-lateralized and the lateral orbitofrontal gyrus is right-lateralized. Language-related regions including pars opercularis and pars triangularis exhibit leftward asymmetry, consistent with the functional dominance of left hemisphere.

Another widely used macroscopic morphological measure is cortical thickness, which is systematically related to the cytoarchitecture and hierarchical structural organization of the cortex^69, 70^. The neurobiological mechanisms underlying developmental changes in cortical thickness are complex and involve processes such as synaptogenesis, proliferation of dendrites, dendritic spines, axonal sprouting, and vascular development^71, 72^. Cortical thickness is a critical biomarker of the underlying neurophysiology and can be used to disentangle cortical dysmaturation from typical development^73^. The IBA shows that cortical thickness increases rapidly during the first year and more slowly in the second year. The increase in cortical thickness during early brain development is positively associated with intelligence and cognitive skills in later stages of life^74, 75^. The IBA also indicates that cortical thickness changes heterogeneously across the cortex (Fig. 2c), in line with previous studies^76^. The association cortices in the temporal, parietal, and prefrontal lobes are thicker^4^ compared with the primary visual and sensorimotor cortices. This is consistent with the functional development of the infant brain where vision, motor, and sensory systems are more mature than executive functions. The IBA also reflects the inter-hemispheric differences in regional cortical thickness: the functionally dominant hemisphere is thinner than the non-dominant counterpart. For instance, the left pars opercularis is thinner, consistent with the functional dominance of the left hemisphere in language production and phonological processing. In contrast, the right pars triangularis is thinner, although the left hemisphere is functionally dominant. This is because pars triangularis is involved in semantic processing that is still imprecise during the first two postnatal years; therefore, the functionally dominant left hemisphere is thicker. The superior temporal gyrus is involved in auditory processing that is pertinent to language acquisition and is dependent on the left hemisphere. The left superior temporal gyrus in IBA is thinner, in line with the early development of infants ability to process audio and spoken language during the study period. Visuospatial processing controlled by the supramarginal gyrus, inferior parietal cortex, and precuneus is dominant in right hemisphere. However, infants do not fully acquire visuospatial processing ability during the first two postnatal years; therefore, these regions in IBA are still thicker in the right hemisphere.

The IBA also reflects cortical maturation in terms of cortical surface area. According to the radial unit hypothesis^77^, cortical surface area expands during infancy due to the symmetrical proliferation of neural progenitors in the ventricular and subventricular zones^78^. Others speculate that intermediate glia cells, generated in the outer subventricular zone, expand in a fan-like manner, and drive the tangential growth of surface area^79, 80^. Prior studies suggest that surface area undergoes a 2- to 4-fold expansion from infancy through adulthood^81^. Postnatal surface area expansion is differential in nature, following regional differences in dendritic length, dendritic spine density, synaptic architecture, and intra-cortical myelination^72, 81–83^. The overexpansion of cortical surface area during early infancy is implicated in disrupted emergence and refinement of cognitive and behavioral skills, and developmental, psychiatric, and neurologic disorders^79, 84, 85^. The IBA reveals that cortical surface area increases by 93% during the first two years after birth (Fig. 4). Findings from our regional analysis are consistent with existing studies^68, 81^: regions with greatest expansion lie in the dorsolateral and medial prefrontal association cortex, lateral temporal, and parietal association cortices, whereas regions with least expansion lie in the occipital cortex, medial temporal cortex, and insula. Areas expanding slowly in the primary and secondary sensory cortices in IBA-S are generally thinner, including the pericalcarine, cuneus, precentral, and postcentral gyri. All regions in IBA-S exhibit significant lateralization. In line with Remer et al.^68^, we found left-lateralization of the medial orbitofrontal gyrus, pars opercularis, superior frontal gyrus, and transverse temporal gyrus and right-lateralization of the superior and middle temporal gyri, inferior parietal cortex, precuneus, frontal pole, paracentral gyrus, pericalcarine, pars orbitalis, and pars triangularis.

In addition to folding geometry, we quantitatively assessed brain tissue volume changes during infancy. Early neurodevelopment is underpinned by cellular and molecular processes that drive the growth and maturation of brain tissues. Cell bodies, axon terminal branches, dendrites, and spines residing in GM contribute to the early growth of GM^86^. Nerve fiber tracts or axons accounting for WM form inter-hemispheric, cortico-cortical, limbic, brainstem, and cortico-spinal connections, providing an efficient network of structural connectivity^87, 88^. Furthermore, recent discoveries reveal that CSF, apart from cushioning the brain, contributes to brain maturation and function by delivering growth factors and signaling molecules to progenitor cells that proliferate into immature neurons, which later migrate to different areas of the cerebral cortex^89, 90^. Age-related brain tissue volume changes are associated with domain-specific cognition^91^ and deviations from typical growth pattern help explain the underlying neurologic impairments, including autism spectrum disorder^92^, ADHD^93^, and schizophrenia^94^. IBA growth curves for GM, WM, ventricular CSF, and whole brain volume (Fig. 4b) indicate that volumes of all tissues increase from birth through two years of age, albeit at rates different from previous studies. Knickmeyer et al.^6^ reported dramatic changes in GM volume by 149% in the first year and 14% in the second year. More moderate changes were reported for the WM volume at 11% in the first year and 19% in the second year. On the other hand, the IBA increases in GM volume by 97.2% and 13.8%, and increase in WM volume by 92.9% and 15.9%, during first and second postnatal years. These differences can be attributed to the cross-sectional study design, different image acquisition protocols, tissue segmentation method that overestimated cortical GM, and the use of ANOVA/ANCOVA for percentage change determination. Only cortical WM is reported by Knickmeyer et al.^6^, resulting in significantly different WM growth rates.

Cortical myelination through infancy was analyzed via T1w/T2w ratio. We observed spatio-temporal changes in myelination of the cerebral cortex throughout the study period, in line with the literature^15^. Highly myelinated regions process information faster than less myelinated regions. The primary visual, motor, and somatosensory cortices are myelinated earlier than the association cortices of the frontal, parietal, and temporal lobes that are involved in higher-order functions, consistent with the functional development of the brain. We also observe a correlation between cortical thickness and myelin content: thicker cortices are lightly myelinated compared to thinner cortices that are heavily myelinated. For instance, the prefrontal cortex has high cortical thickness and low myelin.

Overall, qualitative and quantitative analyses confirm that the surface-volume consistent IBA faithfully reflects the growth trajectories of infants with rich anatomical details. These atlases capture monthly changes in brain shape and size, cortical geometry, tissue contrast, volume, and microstructural characteristics of typically-developing infants. A unique feature of IBA is that its cortical folds are anatomically realistic in both surface and volumetric spaces, as reflected by the good alignment of the cortical surface atlases with tissue interfaces. Another trait that sets the IBA apart from currently available atlases is that it covers each month during the first two postnatal years, providing a dense spatio-temporal depiction of this critical period of development. Note that our analysis currently does not cover the cerebellum and subcortical structures, as precise delineation of cerebellar GM and WM and parcellation of subcortical structures in infants are technically challenging. When proper computational tools become available, our atlases can be employed in the future to investigate cerebellar and subcortical growth.

Discovering irregularities in brain growth at early stages ensures that specialized interventions are accessible to children to optimize long-term health outcomes^95^. We envision that our infant brain atlases will serve a critical role in understanding multi-faceted maturation and identifying aberrant growth. Integrating the cortical surfaces and volumetric atlases in a common coordinate framework will facilitate the development of neuroimaging-based spatio-temporal growth charts—a tool key to investigating regional brain development over time. Our surface-volume atlases will provide a common coordinate space for conflating information from multiple imaging modalities (e.g., structural, diffusion, and functional MRI) for consistent analyses of brain structure and function. Utilizing our temporally dense set of atlases that faithfully capture the month-to-month evolution of the infant brain will allow the precise mapping of anatomical growth patterns from birth through infancy.

In summary, we have delivered for the first time a set of consistent surface-volume atlases developed at each month for the first two years after birth. Collectively, these spatially and temporally dense atlases make an invaluable resource to unravel the early development of brain from macroscale to microscale. Our unique collection of neuroanatomical atlases will equip the neuroscience community with the common reference framework, enabling concurrent surface-volume growth studies. We envisage that conjoint growth analyses will shift the current infant brain development research paradigm and will give new insights into developmental processes underpinning child cognition and social behavior.

## Methods

### Dataset and preprocessing

Longitudinal T1w and T2w MRI scans were acquired using the 3T Siemens Prisma MR scanner with a Siemens 32-channel head coil for 37 subjects enrolled as part of the UNC/UMN Baby Connectome Project (BCP)^32^. These subjects were divided into six cohorts (A_1_, A_2_, A_3_, B_1_, B_2_, B_3_), and each cohort’s first visit was scheduled at 2 weeks, 1, 2, 9, 10, and 11 months, respectively. The subjects in A_1_, A_2_ and A_3_ were scheduled to be scanned every three months in the first year and then at 24 months, whereas the subjects in B_1_, B_2_ and B_3_ were scanned every three months for the first two years. The total number of scans for each subject is different since all subjects cannot be scanned at all expected time points. A total of 108 scans were used in the current work.

T1w MR images were acquired with 208 sagittal slices, TR/TE = 2400/2.24 ms, flip angle = 8, acquisition matrix = 320 × 320, and resolution = (0.8 mm)^3^. T2w MR images were acquired with 208 sagittal slices, TR/TE = 3200/564 ms, variable flip angle, acquisition matrix = 320 × 320, and resolution = (0.8 mm)^3^. The study protocols were approved by the Institutional Review Board of the School of Medicine of The University of North Carolina at Chapel Hill (UNC-CH), NC, USA.

All MR images were preprocessed using our infant-centric processing pipeline consisting of the following steps: (i) rigid alignment of T1w and T2w images using FLIRT^96^; (ii) skull stripping by a learning-based method^97^; (iii) intensity inhomogeneity correction by N3^98^; (iv) brain tissue segmentation by an infant dedicated learning-based method^99^; (v) hemisphere separation and subcortical filling; and (vi) topologically-correct cortical-surface reconstruction^100, 101^.

### Surface-volume atlas construction

Our atlas construction method (Supplementary Fig. 20a) involves two main tasks: (i) construction of 12-month surface-volume atlas via surface-constrained groupwise registration; and (ii) longitudinal atlas construction for 2 weeks to 24 months by propagating the 12-month atlas through parallel transported longitudinal deformations. The steps involved in each task are detailed below.

### Construction of reference atlas — IBA12

We first construct the 12-month surface-volume reference atlas, i.e., IBA12, and use it to generate the atlases at the other time points. The IBA12 lies in the middle of the two-year time span and is hence ideal to capture developmental patterns between birth and two years of age. In the BCP dataset, the 12-month time point has the greatest number of scans.

#### Surface-constrained groupwise registration

Infant brain MRI exhibits dynamic structural changes from birth to 2 years of age. The brain undergoes remarkable changes in size, shape, and tissue contrast. The low intensity contrast of infant brain MRI renders image registration and atlas construction challenging. The combined volumetric and surface (CVS) registration method^102^ concurrently aligns volumetric tissue boundaries along with the white and pial cortical surfaces. However, CVS registration caters mainly to adult brain images and is not designed specifically for infant brain MRI. Rapid appearance changes affect both segmentation and registration accuracy. Here, we use a dynamic elasticity model with surface constraint (SC-DEM^33^) for groupwise registration using the tissue segmentation maps instead of the intensity images. Groupwise registration allows a population of tissue segmentation maps to be registered simultaneously to a common space as demonstrated in our previous work^31^. In our case, this is achieved by first selecting a reference based on a subject scan that is most similar in appearance to the whole set of images. Then, the reference tissue segmentation map is iteratively updated by fusing all tissue segmentation maps that are registered to it. SC-DEM improves the alignment of tissue boundaries using predetermined surface transformations. Before SC-DEM groupwise registration, the tissue segmentation maps are affine-aligned with the reference tissue segmentation map using FSL FLIRT^103^. For simplicity, in the rest of the paper affine transform is assumed where appropriate and will not be explicitly stated.

For cortical surface registration, the white surface is inflated and mapped onto the unit sphere via metric distortion minimization^38^. The surfaces in spherical space are represented by two folding attributes: average convexity and mean curvature. The spherical surfaces of the moving scans are registered to that of the reference scan using Spherical Demons^34^. The resulting vertex-wise correspondences are propagated to the white and pial cortical surfaces by leveraging the one-to-one vertex mapping between the spherical surfaces, and the white and pial cortical surfaces.

SC-DEM employs the dynamic elasticity model (DEM^104^) to characterize large nonlinear displacements. DEM registration is formulated based on the principle of elastodynamics. An image is modeled as an elastic body that is displaced from its resting state by an external force. This disturbance propagates across the whole body as elastic waves, representing the nonlinear displacements. For the *i*-th subject scanned at time^5^ *τ* ∈ 𝒲_12_, where 𝒲_*ρ*_ = [(*ρ* − 1.5), (*ρ* + 1.5)] months, we estimate displacement field 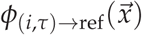 by solving the wave equation:

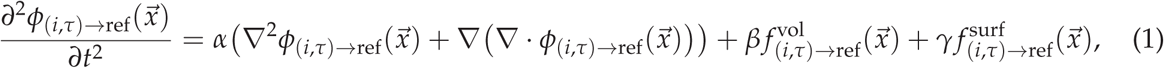

where ∇^2^ is the vector Laplacian operator and 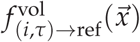 and 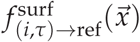 are respectively the volumetric and surface force fields. The volumetric force field 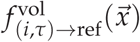 captures the misalignment between the warped moving tissue segmentation map 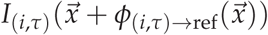 and the reference tissue segmentation map 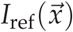:

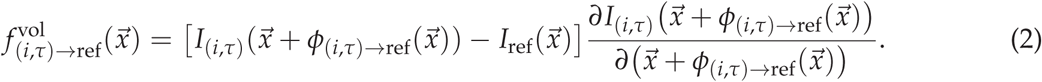

The surface force field 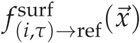 is computed based on the differences between the volumetric displacements and the predetermined surface displacements for white and pial surfaces for both left and right hemispheres. *α* is a parameter that controls deformation smoothness. *β* and *γ* balance the two force fields and control the rate of convergence. Registration halts when the two force fields are negligibly small, thereby consistently aligning the surface and the tissue segmentation map.

#### Cortical surface atlas

The cortical surface atlas for white and pial surfaces of both left and right hemispheres is constructed by using the registered cortical surfaces 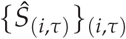, each associated with a weight *w*(*τ*, 12), computed using 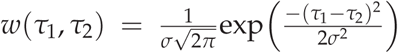, where parameter *σ* controls temporal smoothness and is set to 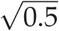 month. The 12-month cortical surface atlas 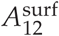 is computed by weighted averaging of the vertex coordinates for 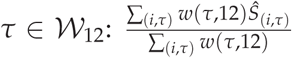. Next, we use 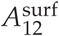 to construct the corresponding volumetric atlas such that the two atlases are defined in the same coordinate space.

#### Volumetric atlas

The tissue segmentation map based volumetric atlas 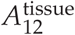 is constructed by correcting the displacement fields 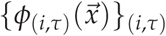 for *τ* ∈ 𝒲_12_ based on the surface atlas 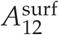, ensuring the alignment of the volumetric WM-GM and GM-CSF interfaces in correspondence to the surface atlas. This involves updating the volumetric displacement fields by considering the surface misalignments as described below:

i. Compute for each vertex with coordinates 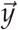 on the surface atlas the vertex-wise surface displacement 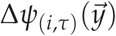 between *Ŝ*(*i,τ*) and 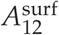.
ii. For each query voxel 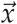, find in 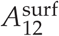 the triangle with vertices 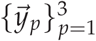 that is closest to the query voxel.
iii. Compute the corrective volumetric displacement field 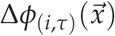 as

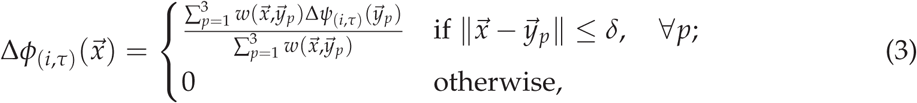

where 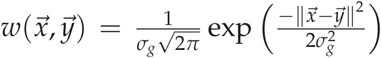 and *δ* = 6 mm. We set *σ*_*g*_ = 3 mm for smooth displacement fields.
iv. Warp the tissue segmentation maps using the total displacement field 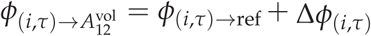 for better alignment and preservation of anatomical details.
v. Determine the tissue label at location 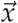 via majority voting for *τ* ∈ 𝒲_12_:

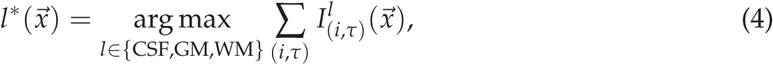

where

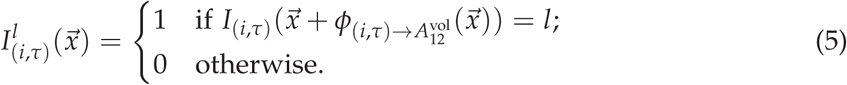
vi. Warp the intensity images (T1w and T2w) using the total displacement field 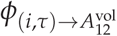 to obtain 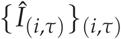.
vii. Average the warped intensity images using the weights *w*(*τ*, 12) to obtain 12-month T1w and T2w atlases: 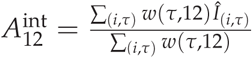.

### Construction of longitudinal surface-volume atlases

The surface-volume atlases for 2 weeks to 24 months are generated by propagating the IBA12 to each month. To achieve this, we determine the average longitudinal change from each month to the 12-month atlas space. This is realized by transporting the longitudinal deformations to the atlas space via inter-subject deformations (Supplementary Fig. 20b). Next, we detail our method for age-dependent atlas construction.

#### Parallel transport of longitudinal deformations

The longitudinal growth of a subject is characterized by the displacement field 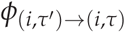 from time point *τ*^*′*^ to time point *τ* ∈ W_12_, estimated via affine and SC-DEM registration. Intra-subject deformation fields 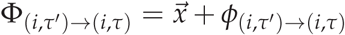 are spatially normalized to the 12-month atlas space by parallel transport^105–107^ — disentangling — inter-subject variability from longitudinal growth. The process involves warping and reorientation of the deformations from the subject space to the 12-month atlas space. The transported deformation field 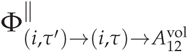 at location 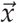 is given as 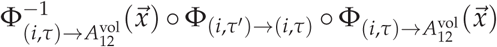.

#### Kernel regression

To construct age-specifc surface-volume consistent atlases, we warp the 12-month atlas to each specific time point *τ*_0_ with weighted average of the transported displacement fields 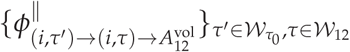

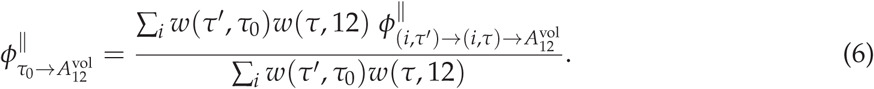

The transported displacement fields are weighted based on whether *τ*^*′*^ and *τ* are close to *τ*_0_ and 12, respectively (Supplementary Fig. 20c). The displacement field 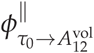 is used to warp the 12-month cortical surface and tissue segmentation maps atlases to each time point. Intensity images (T1w and T2w) are warped with total deformation fields 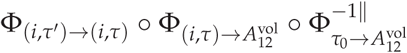 to obtain 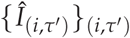 and then averaged to generate T1w and T2w atlases: 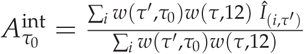.

### Developmental trajectories of the infant population

In order to characterize infant brain maturation, we estimated the developmental trajectories of brain tissues volume, average convexity, mean curvature, cortical thickness, surface area, and cortical myelination. For subject *i* at scan time *t*, the age effects on feature *f*_*i*_(*t*) are modeled using GAMM^39^: *f*_*i*_(*t*) = *s*(*t*) + *γ*_*i*_. Here, *s*(.) is a smooth nonlinear function fitted by cubic regression splines and *γ*_*i*_ is the subject-specific random intercept. The model internally selects the degree of smoothness using the restricted maximum likelihood (REML) criterion^108^ in R.

### Statistical analysis

We performed one-sample two-tailed *t*-test at 1% significance level to check whether geometric features of the cortical ROIs are significantly lateralized.

### Atlas construction with compared methods

For comparison purposes, we constructed longitudinal cortical surface atlases with Spherical Demons ^34^ and volume and surface atlases with ANTs ^35^.

#### Spherical Demons atlas construction

The cortical surface atlases at each month *τ*_0_ are generated by groupwise registration of white surfaces of the subjects scanned at time 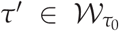. The white cortical surfaces are mapped onto the unite sphere and then spatially aligned via Spherical Demons that uses average convexity and mean curvature to drive surface registration. During groupwise registration, individual cortical surfaces are aligned to the mean cortical surface that is iteratively updated. Finally, average convexity and mean curvature maps are averaged to obtain the surface atlases.

#### ANTs atlas construction

We first generate the 12-month surface and volume atlases 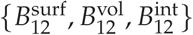, and use these to obtain atlases at the other time points.

For 12-month atlas, the tissue segmentation maps of the subjects scanned at time *τ* ∈ 𝒲_12_ are spatially aligned via groupwise registration with ANTs. The cortical surfaces and intensity images of the registered subjects are then averaged with weights *w*(*τ*, 12). The warped tissue segmentation maps are fused via majority voting. Next, we propagate these 12-month surface and volume atlases to each month *τ*_0_ via parallel transport of the intra-subject deformation fields estimated using affine and ANTs registration for subjects scanned at *τ* and perform kernel regression of the transported displacements 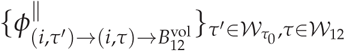 (Eq. 6) to obtain longitudinal atlases. Warped intensity images 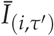 are used to obtain the T1w and T2w atlases: 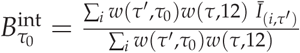.

## Supplementary Information

The manuscript contains supplementary material.

## Acknowledgements

This work was supported in part by the National Institutes of Health (NIH) grants (EB008374, MH125479, MH116225, MH117943, and MH110274) and the efforts of the UNC/UMN Baby Connectome Project Consortium.

## Author Contributions

S.A.: Methodology, Software, Investigation, Writing - Original Draft, Writing – Review & Editing. Y.W.: Writing – Review & Editing. Z.W.: Data Curation. K.-H.T.: Data Curation. W.L.: Resources. G.L.: Resources. L.W: Resources. P.-T.Y.: Conceptualization, Supervision, Funding Acquisition, Validation, Writing – Review & Editing.

## Data Availability

The data used in this paper were provided by the investigative team of the UNC/UMN Baby Connectome Project. The data can be obtained from the National Institute of Mental Health Data Archive (NDA) (http://nda.nih.gov/) or by contacting the investigative team32.

## Correspondence

Correspondence and requests for materials should be addressed to P.-T.Y.

## Footnotes

1 Mapping is only performed for IBA12-S and ANTs. Spherical Demons outputs only spherical atlases of cortical features.

2 Growth rates are reported in terms of percentage change.

3 IBA cortical ROIs are shown in Supplementary Fig. 10

4 Thicker cortices are less matured.

5 Scan time is defined as the chronological age.

