## Supplementary Materials for "Multifaceted Atlases of the Human Brain in its Infancy"

### Supplementary Information

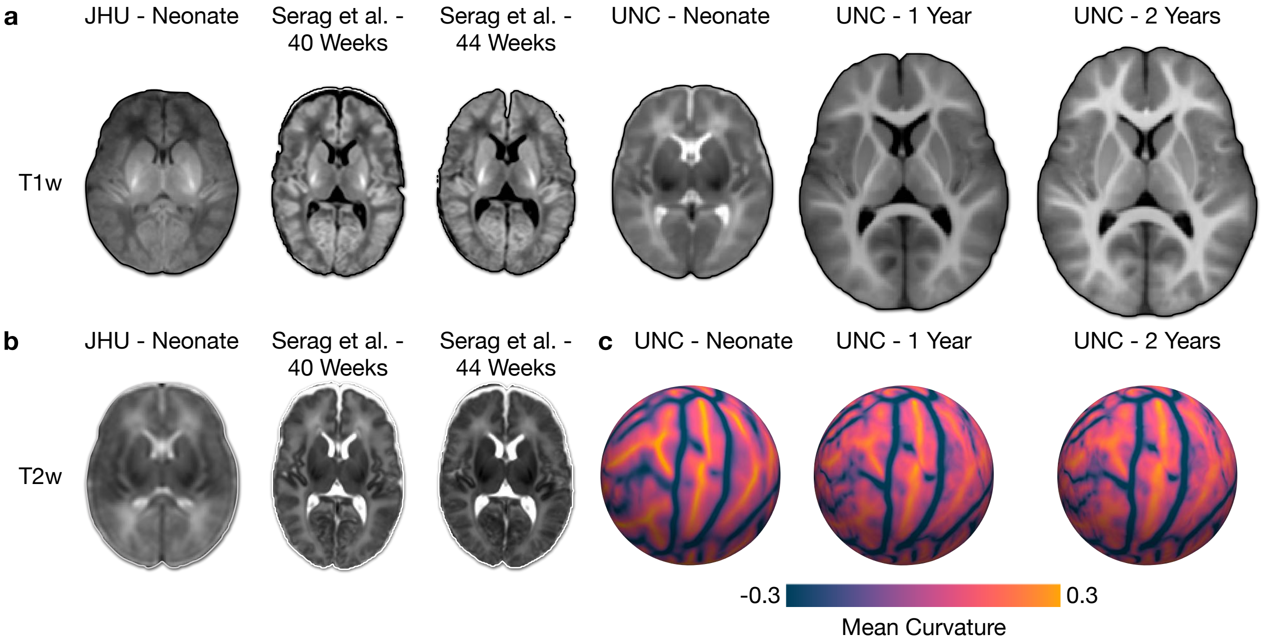


**Supplementary Fig. 1 | Existing neonatal and infant atlases. a**,**b**, Commonly used T1w and T2w atlases representing neonatal and infant populations. **c**, Spherical surface atlases representing mean curvature of neonates and infants.

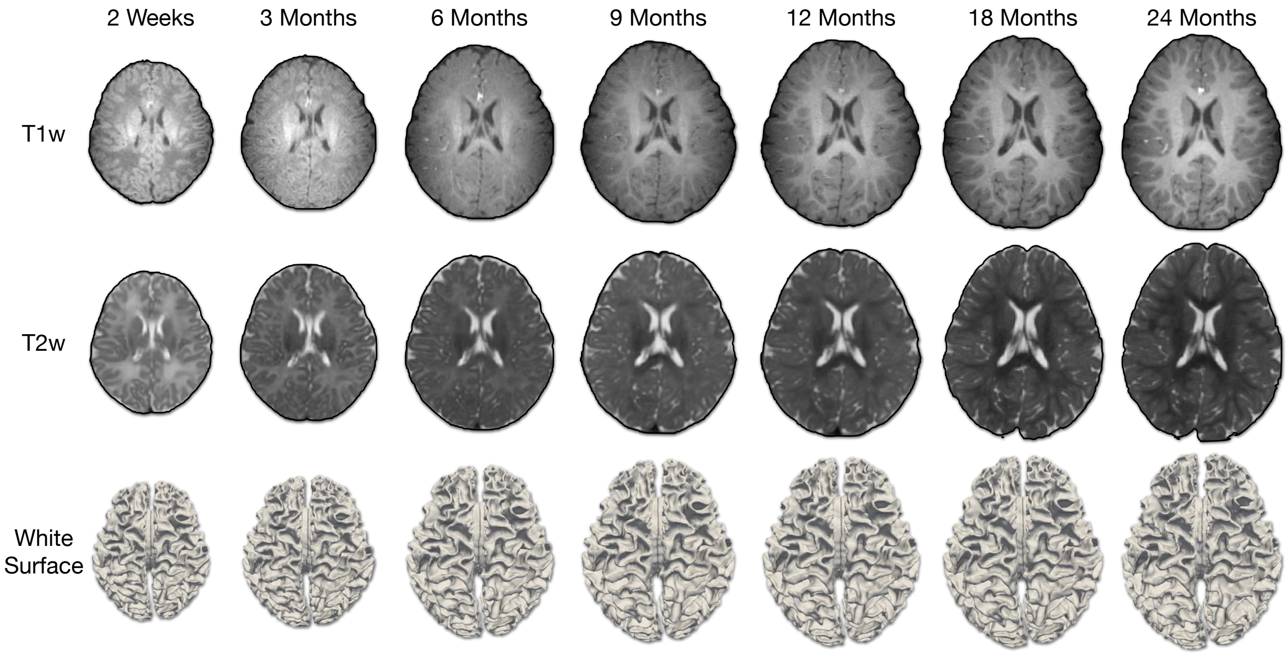


**Supplementary Fig. 2 | Longitudinal infant scans.** T1w and T2w images and white surfaces of an infant scanned between two weeks and two years of age.

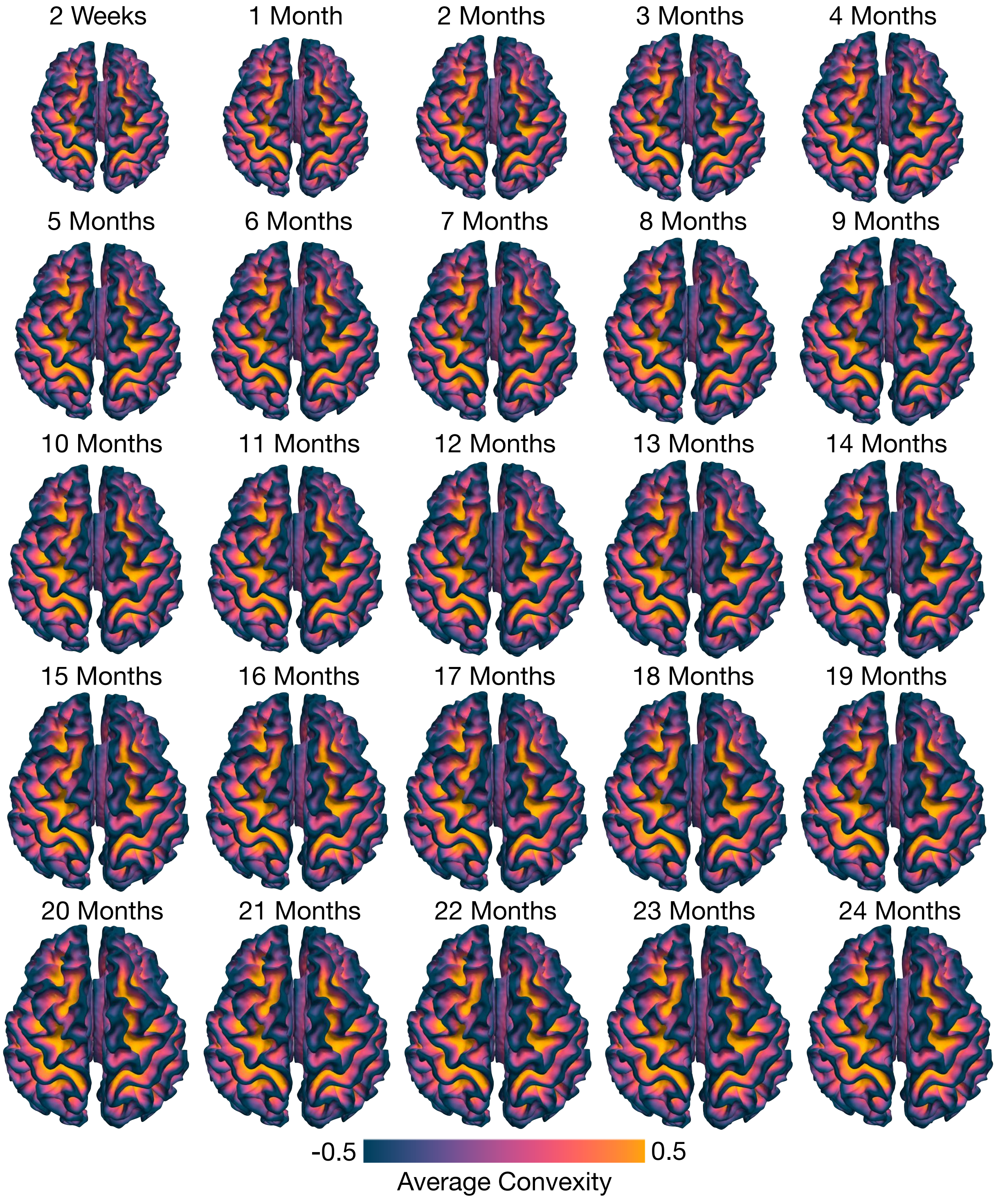


**Supplementary Fig. 3 | Cortical atlases of the white surface.** The age-specific cortical atlases of the white surface for both hemispheres, color-coded by average convexity. The gyri and sulci are present consistently from 2 weeks to 24 months.

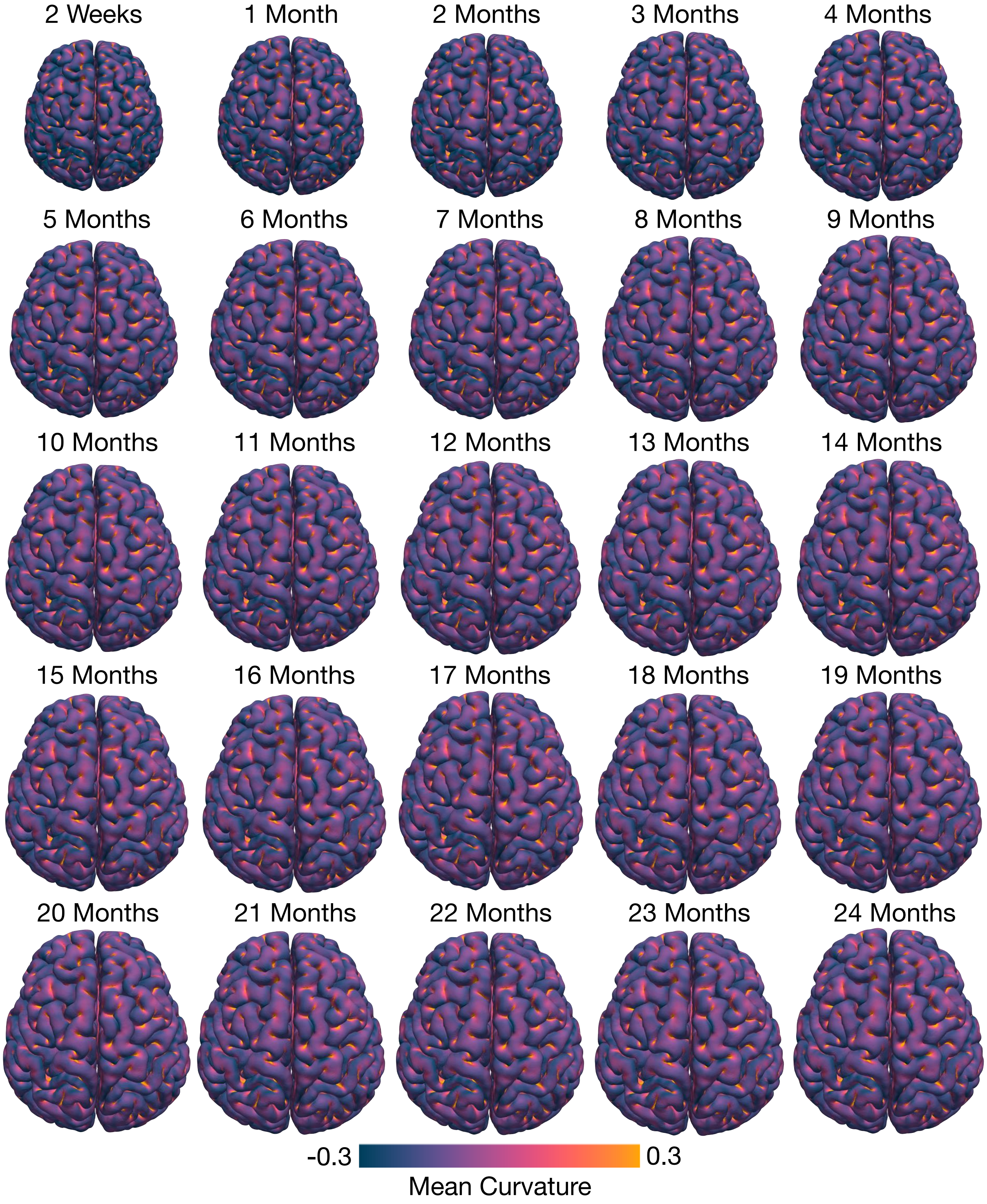


**Supplementary Fig. 4 | Cortical atlases of the pial surface.** Dorsal views of the age-specific cortical atlases of the pial surface for both hemispheres spanning 2 weeks to 24 months, color-coded by mean curvature.

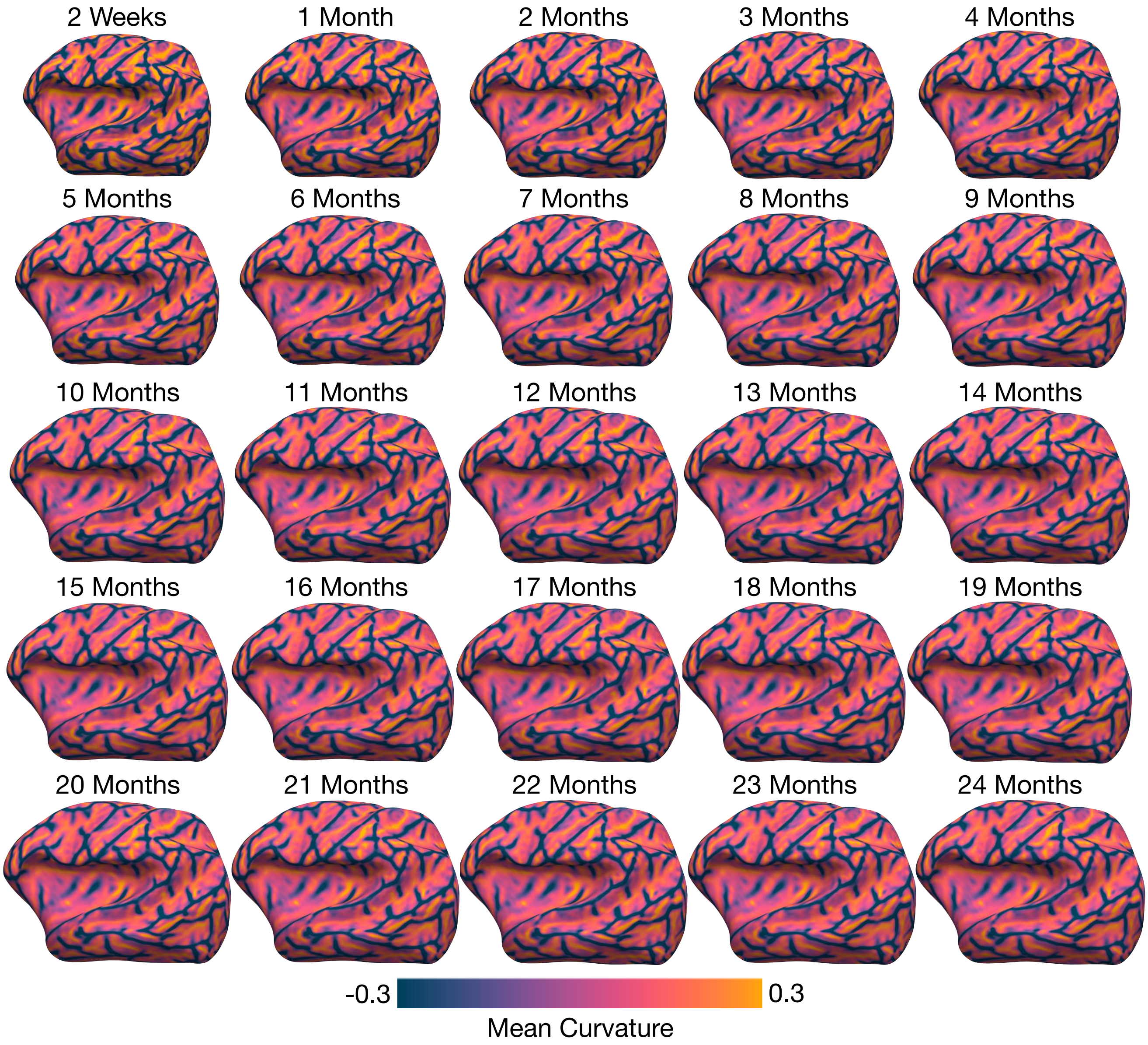


**Supplementary Fig. 5 | Mean curvature maps of the longitudinal infant brain atlases.** The inflated cortical atlases of the white surface (left hemisphere, lateral view) superimposed with mean curvature maps spanning 2 weeks to 24 months.

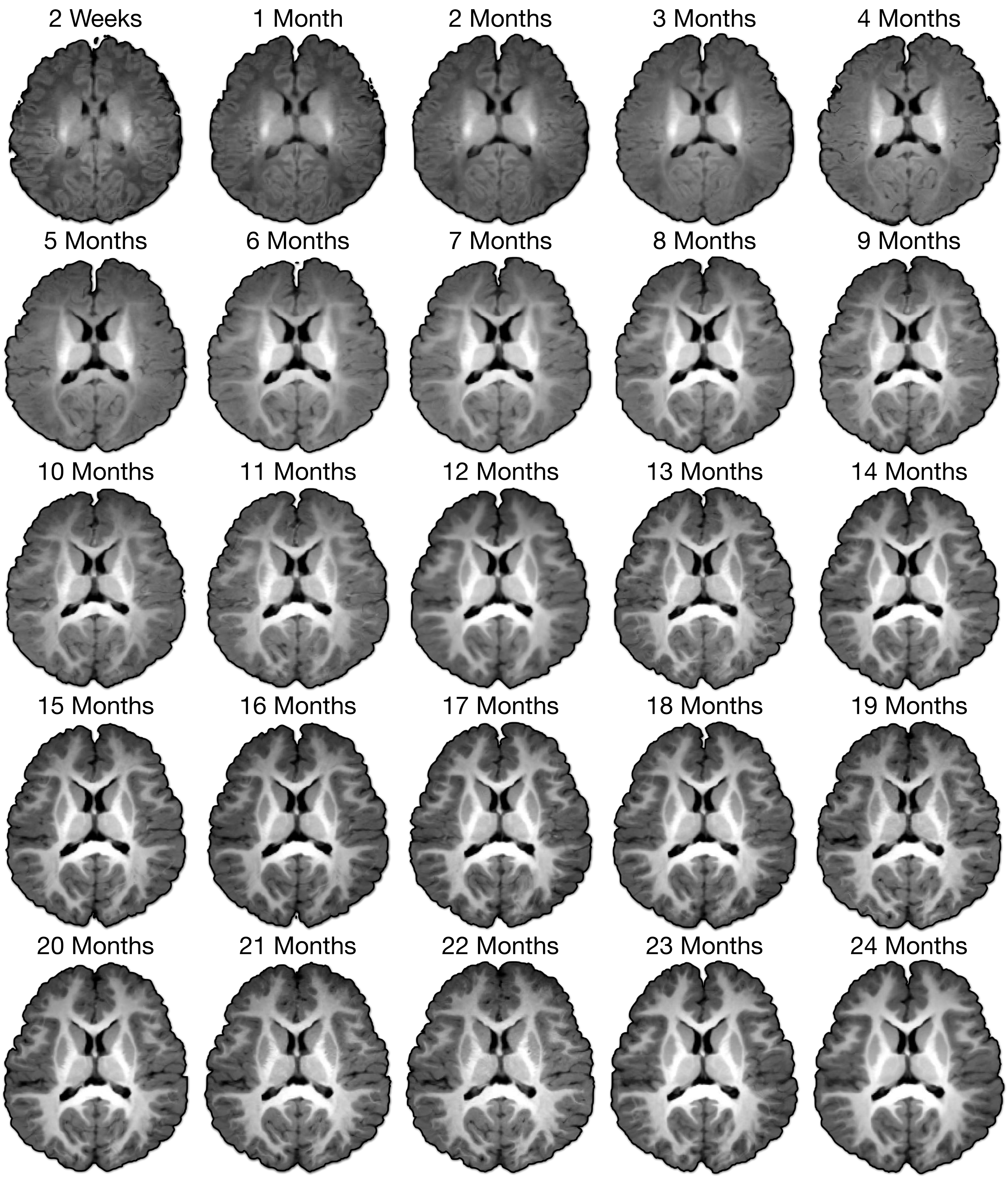


**Supplementary Fig. 6** | **T1w atlases of the infant brain.** Transverse sections of the T1w atlases depicting dynamic changes in tissues contrast, size, and shape of the anatomical structures at each month between 2 weeks and 24 months.

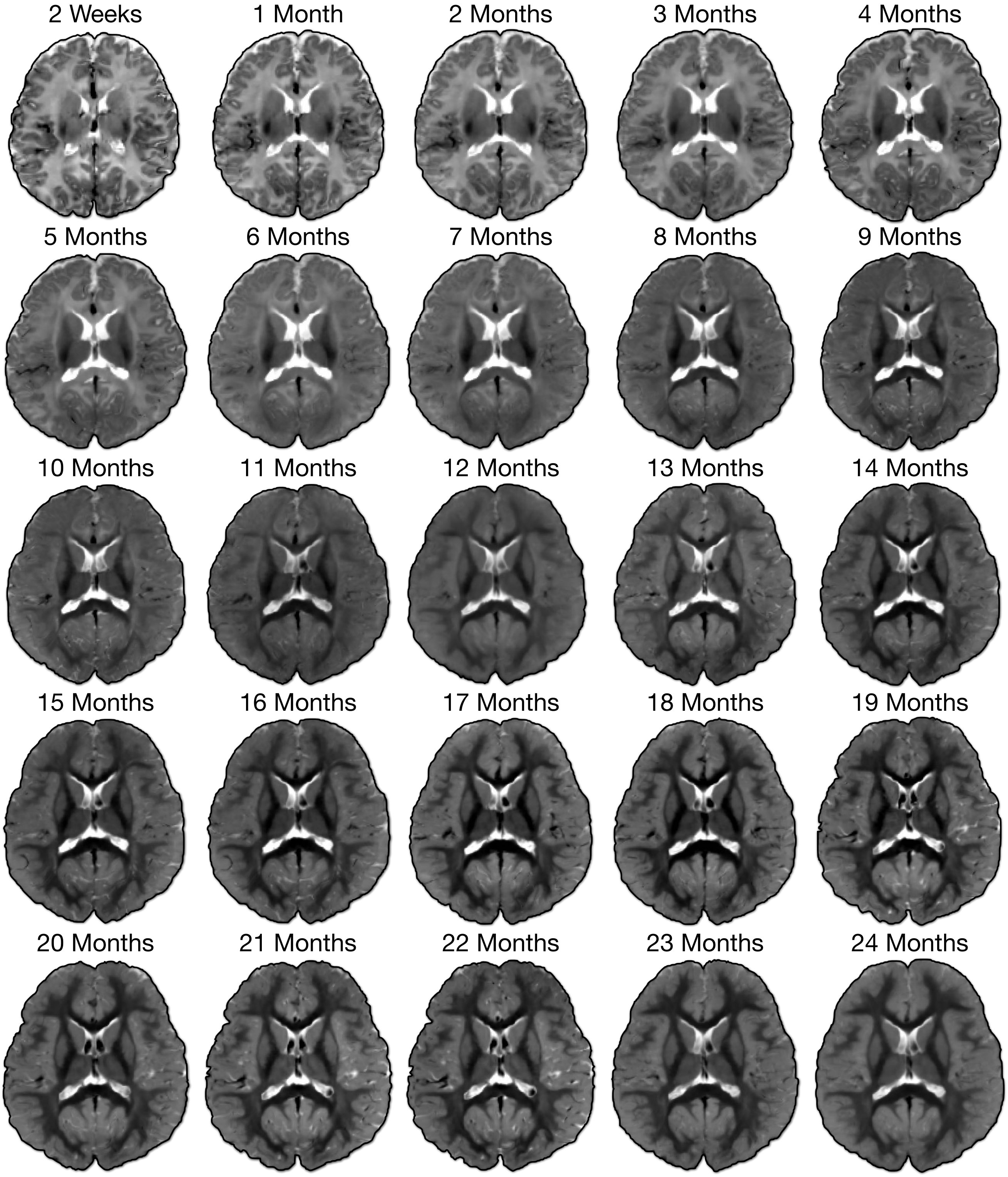


**Supplementary Fig. 7 | T2w atlases of the infant brain.** Transverse sections of the T2w atlases depicting dynamic changes in tissues contrast, size, and shape of the anatomical structures at each month between 2 weeks and 24 months.

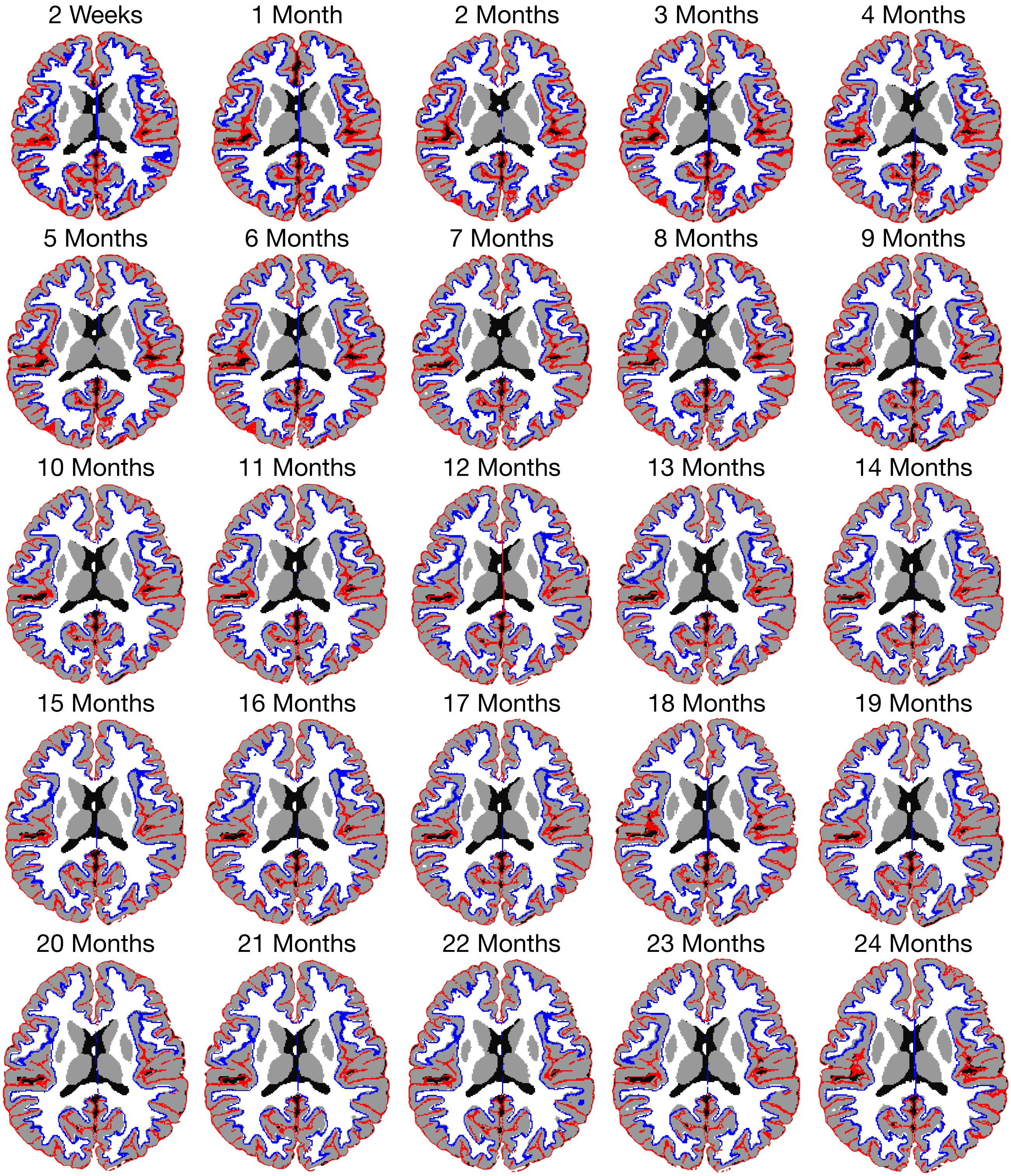


**Supplementary Fig. 8 | Spatial consistency between age-specific cortical surface and volumetric atlases.** Left and right cortical atlases of the white (*blue*) and pial (*red*) surfaces are superimposed onto the tissue map atlases.

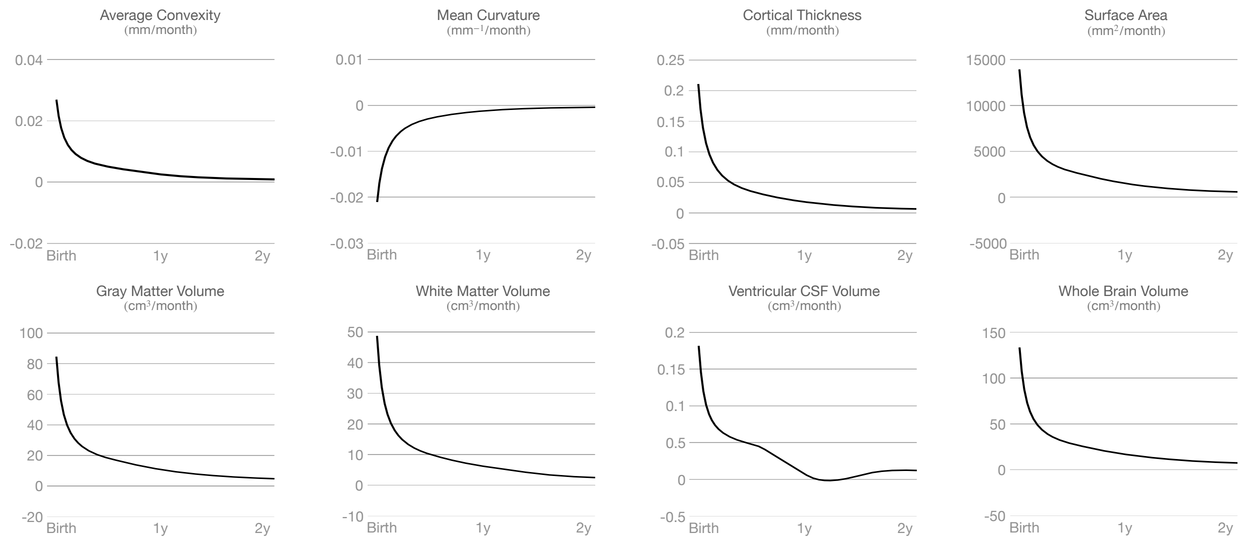


**Supplementary Fig. 9 | Cortical and volumetric developmental velocities.** IBA velocity curves for surface and volumetric features, estimated via the first derivatives of generalized additive model (GAM)-fitted trajectories.

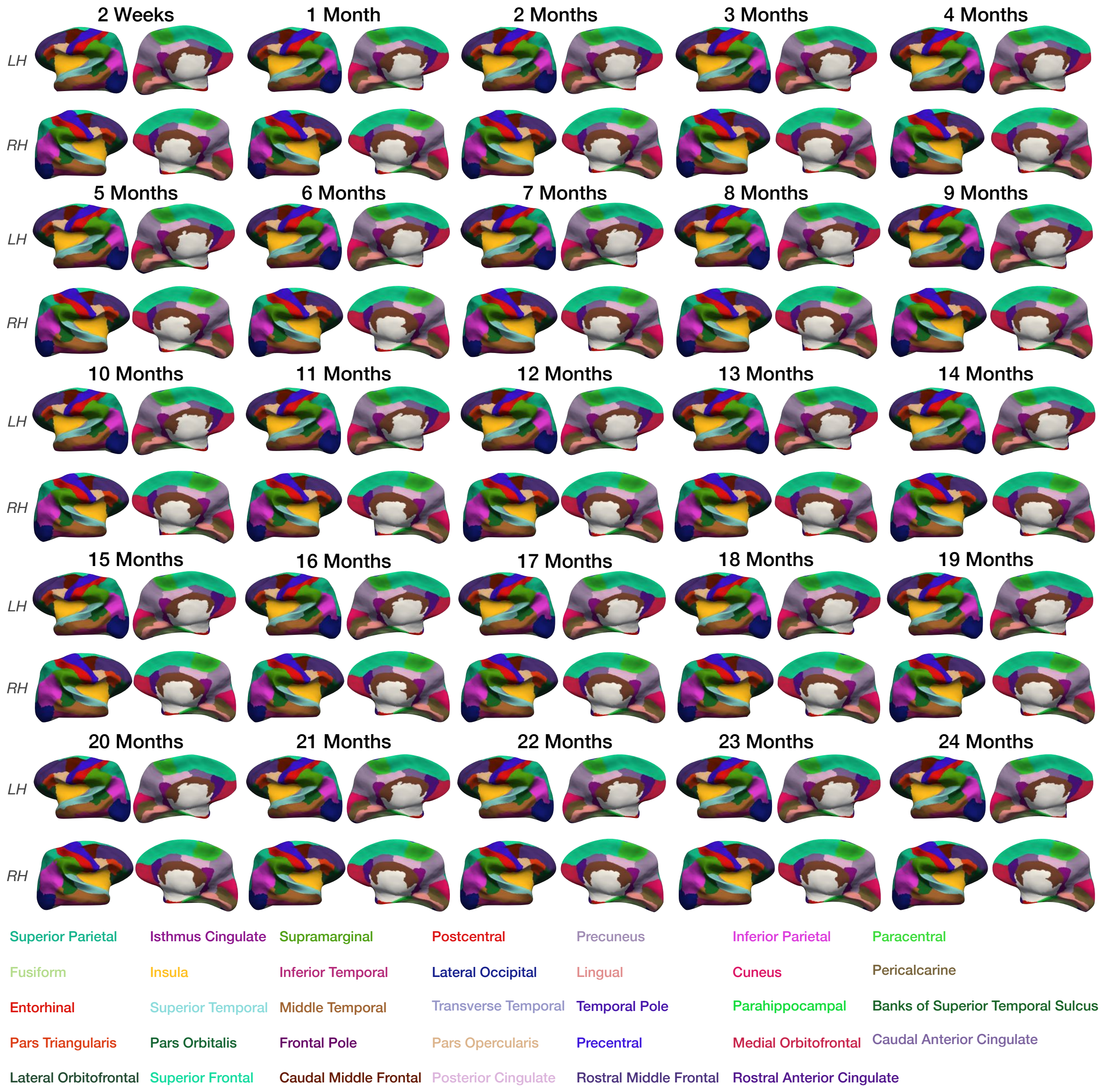


**Supplementary Fig. 10 | Cortical ROIs.** Desikan-Killiany cortical parcellation mapped onto the inflated white surfaces of the IBA.

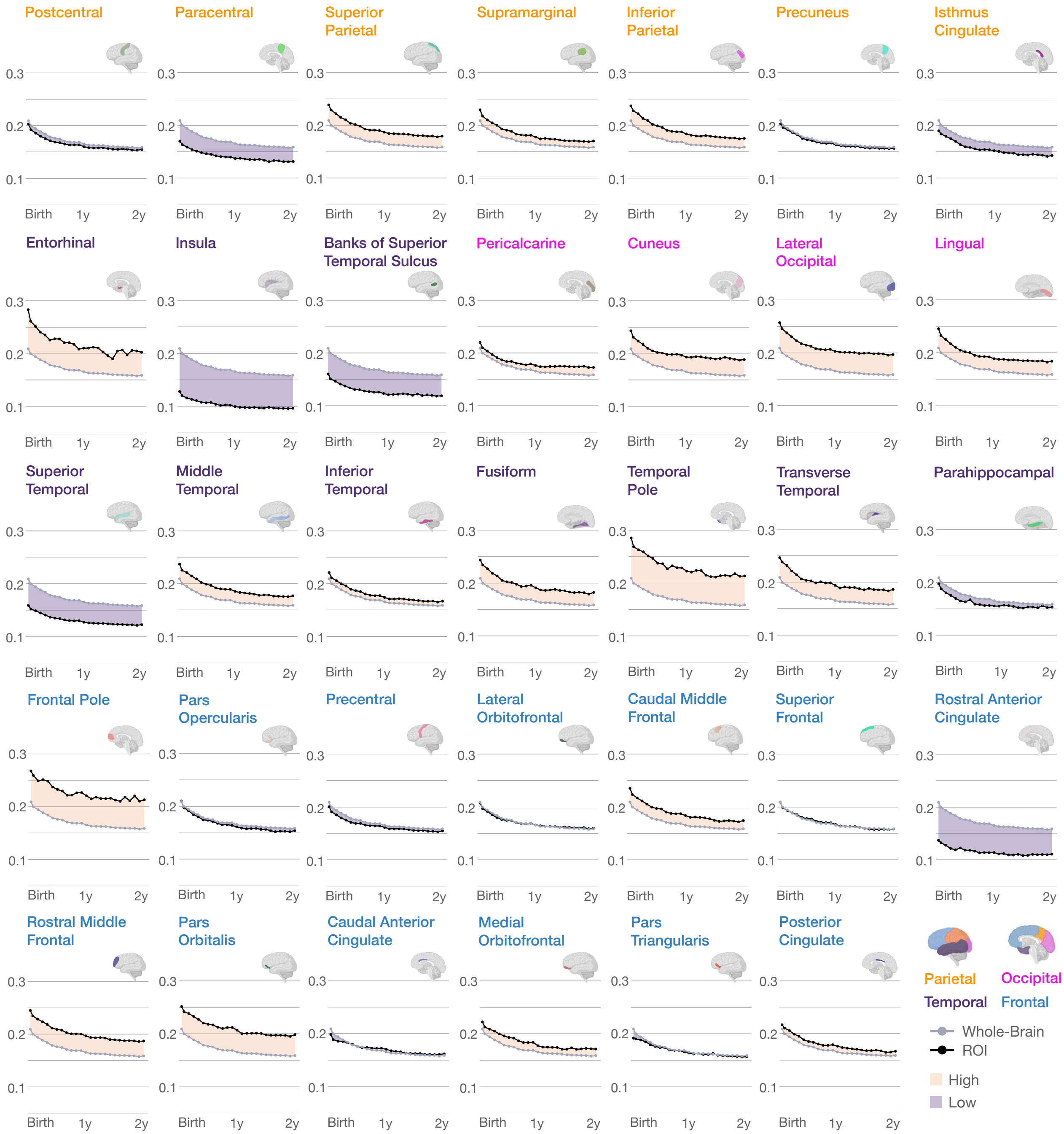


**Supplementary Fig. 11 | Regional developmental trajectories of mean curvature.** Growth curves of mean curvature for IBA cortical regions. Shaded regions indicate whether mean curvature is higher or lower than average.

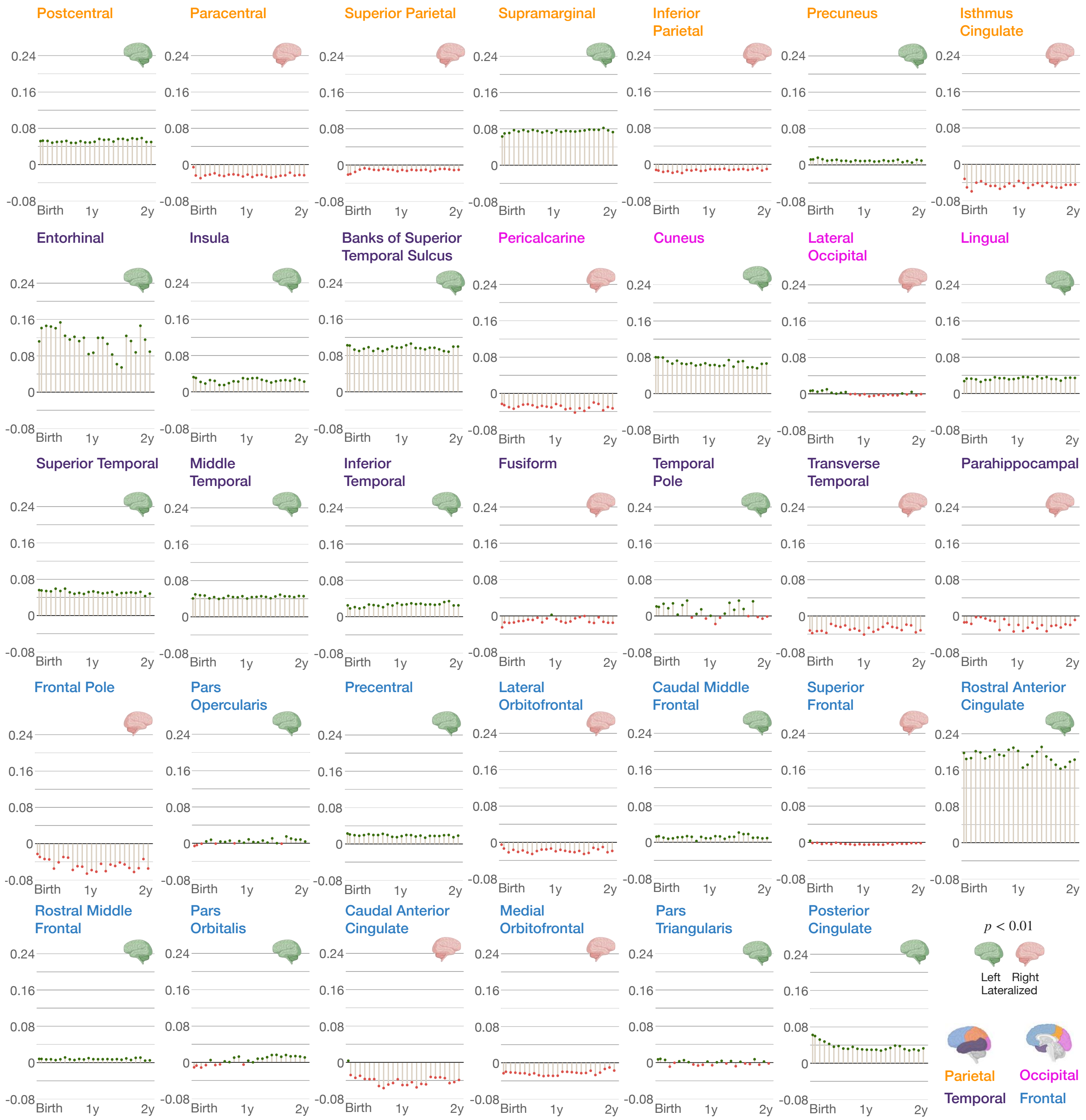


**Supplementary Fig. 12 | Hemispheric asymmetry of mean curvature.** Region-specific laterality index for mean curvature of the IBA. Positive laterality is associated with left lateralization ($p < 0.01$) and negative laterality is associated with right lateralization ($p < 0.01$).

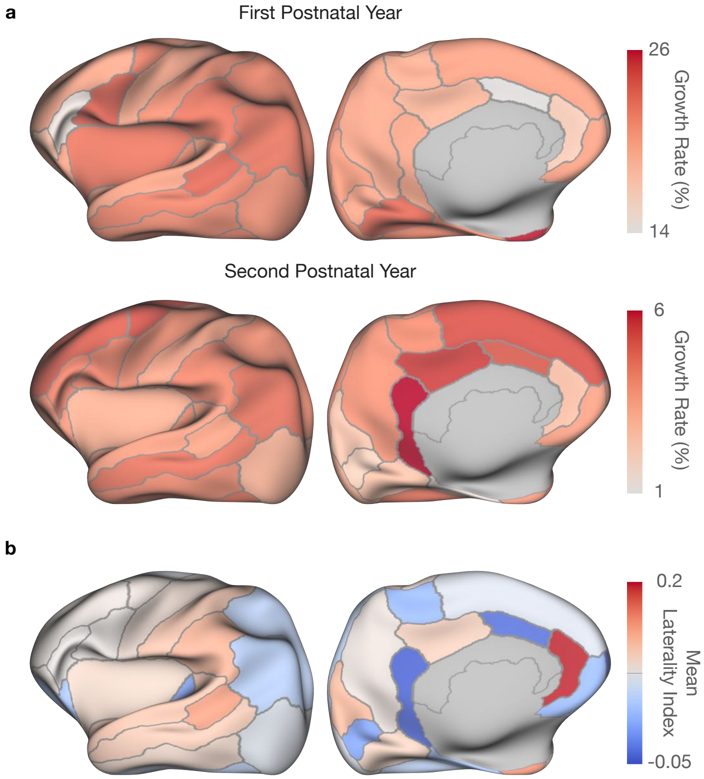


**Supplementary Fig. 13** | **Analysis of mean curvature.** **a**, Regional growth rates in terms of mean curvature for the first (*top row*) and second (*bottom row*) postnatal years. **b**, ROI-specific mean laterality index for mean curvature.

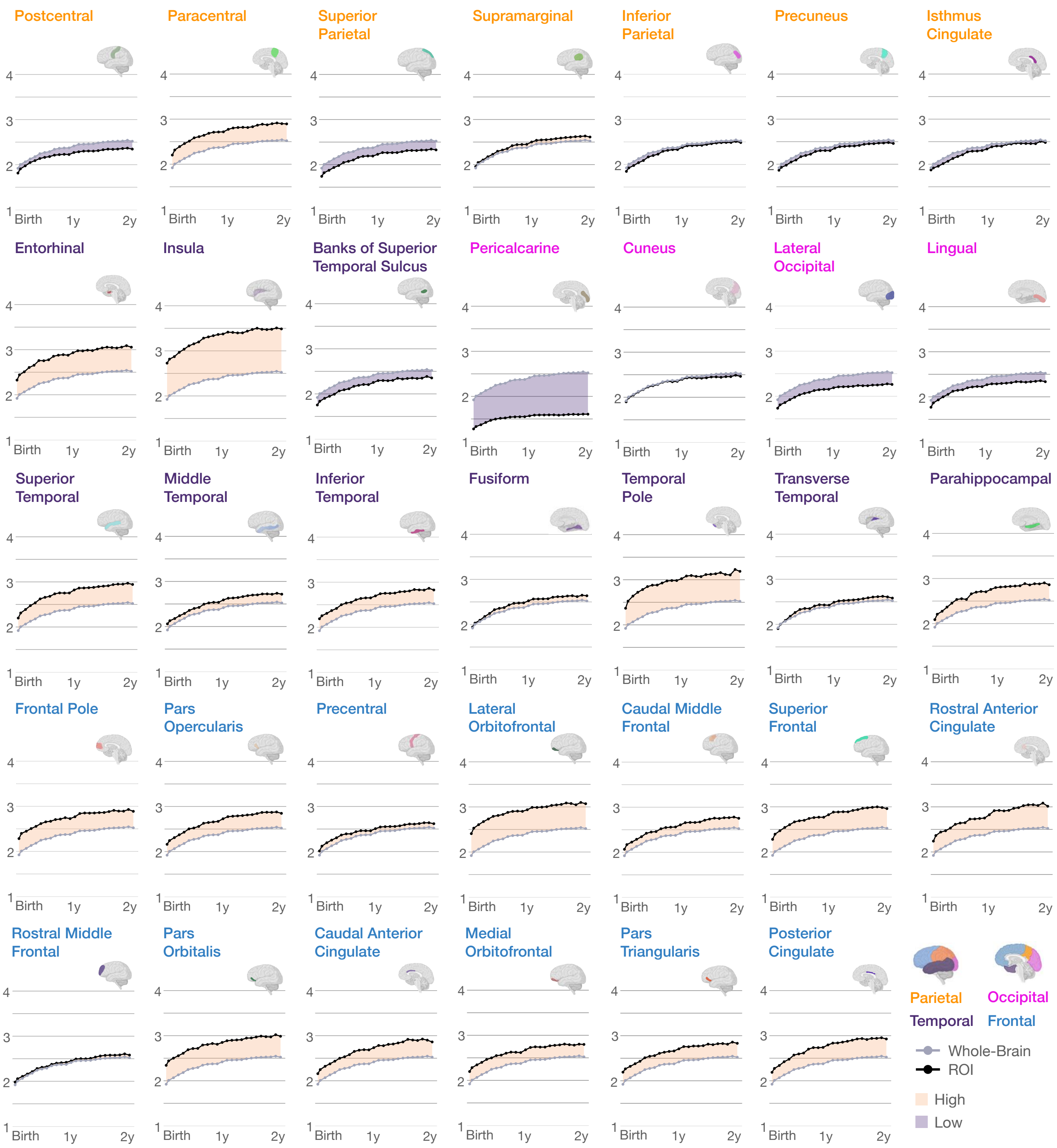


**Supplementary Fig. 14 | Regional developmental trajectories of cortical thickness.** Growth curves of cortical thickness for IBA cortical regions. Shaded regions indicate whether cortical thickness is higher or lower than average.

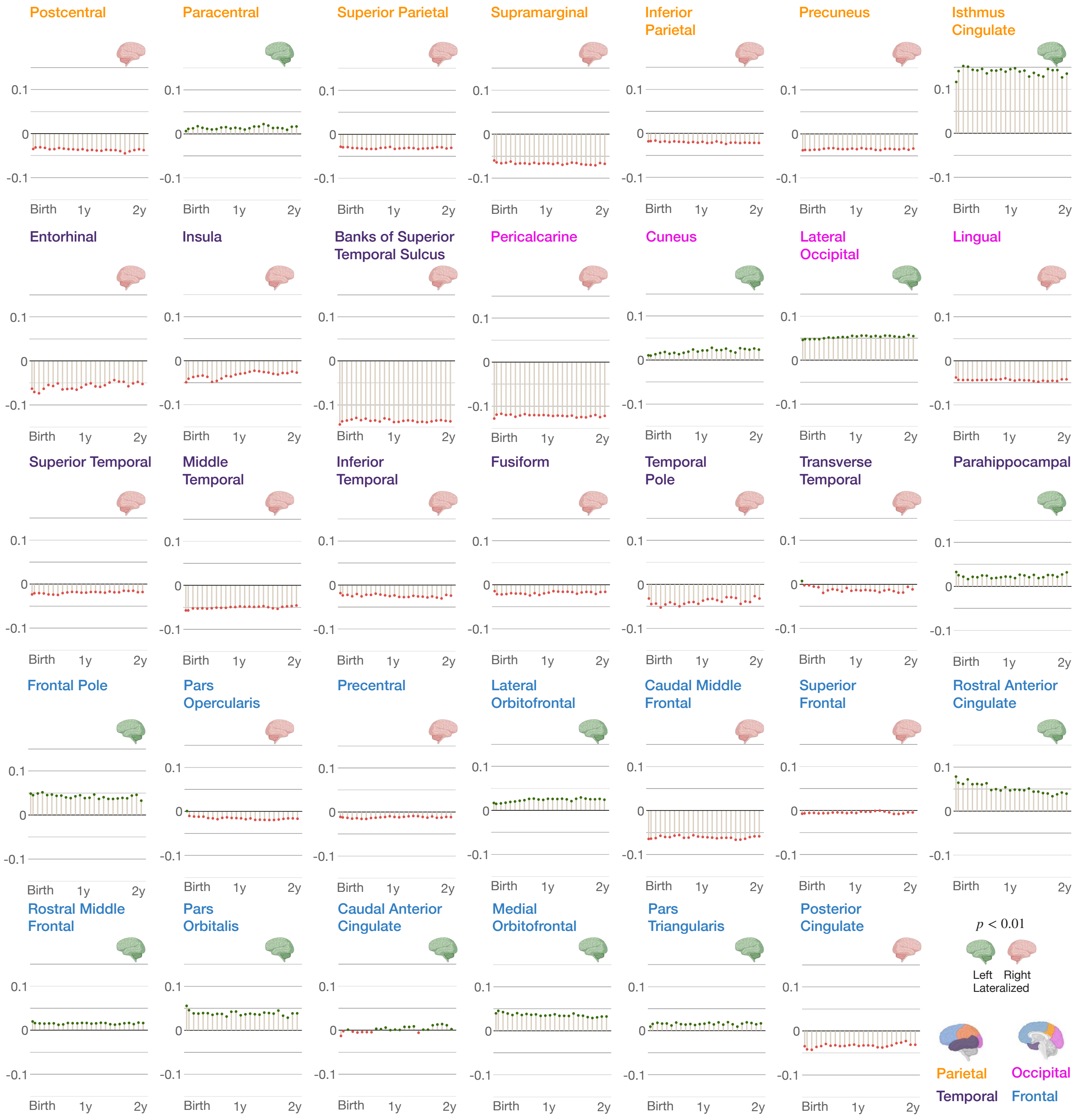


**Supplementary Fig. 15 | Hemispheric asymmetry of cortical thickness.** Region-specific laterality index for cortical thickness of the IBA. Positive laterality is associated with left lateralization ($p < 0.01$) and negative laterality is associated with right lateralization ($p < 0.01$).

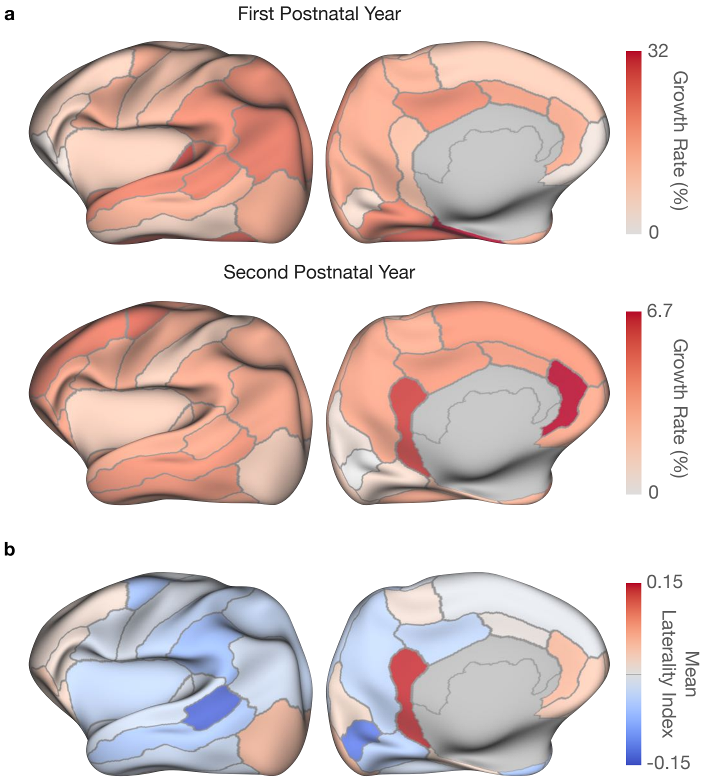


**Supplementary Fig. 16 | Analysis of cortical thickness.** **a**, Regional growth rates in terms of cortical thickness for the first (*top row*) and second (*bottom row*) postnatal years. **b**, ROI-specific mean laterality index for cortical thickness.

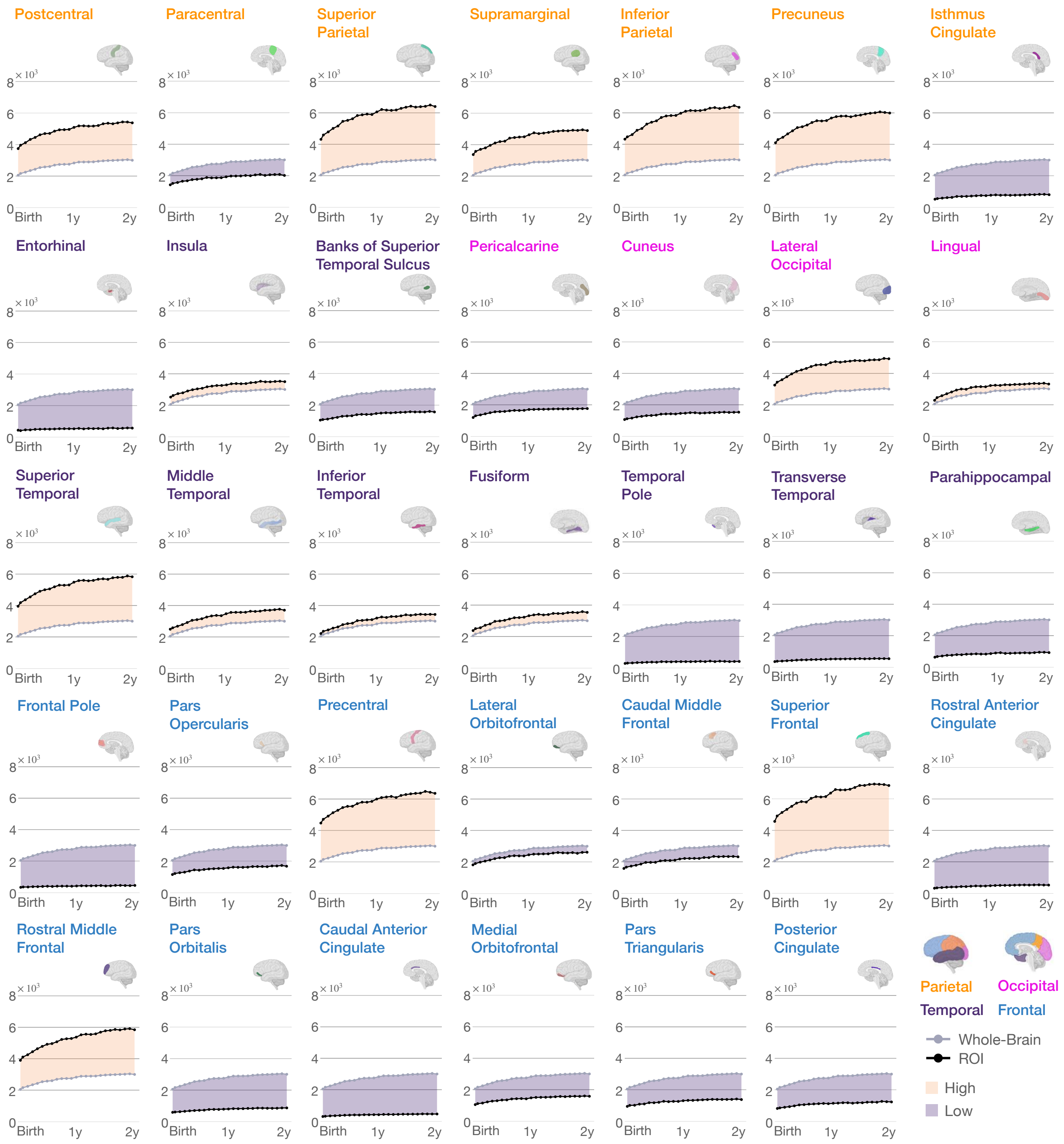


**Supplementary Fig. 17 | Regional developmental trajectories of surface area.** Growth curves of surface area for IBA cortical regions. Shaded regions indicate whether surface area is higher or lower than average.

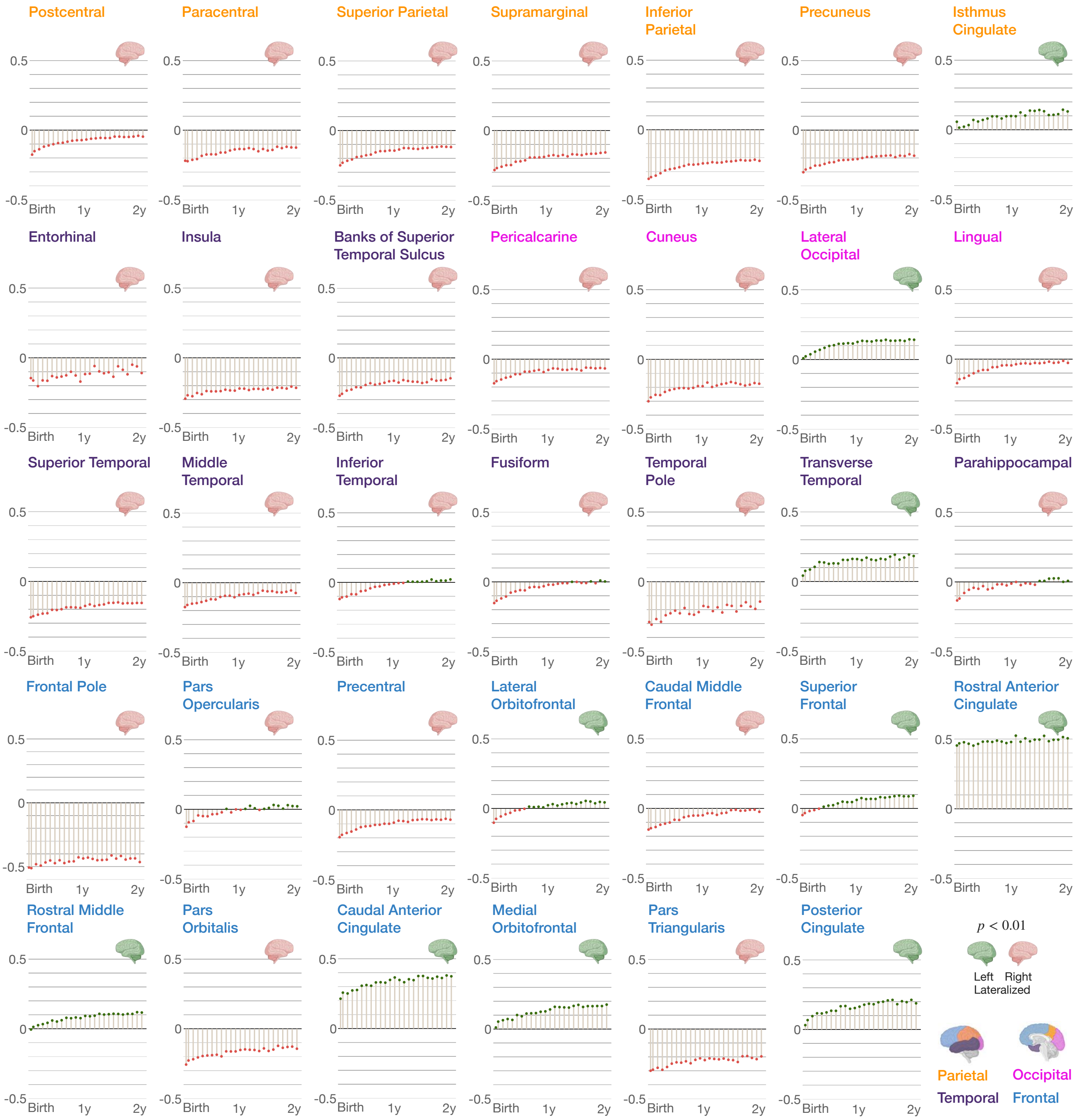


**Supplementary Fig. 18 | Hemispheric asymmetry of surface area.** Region-specific laterality index for surface area of the IBA. Positive laterality is associated with left lateralization ($p < 0.01$) and negative laterality is associated with right lateralization ($p < 0.01$).

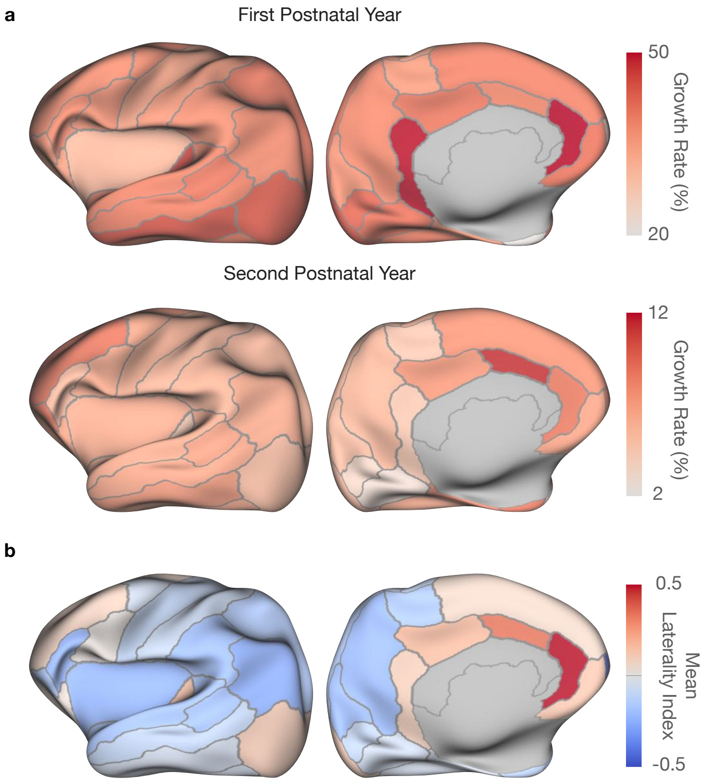


**Supplementary Fig. 19** | **Analysis of surface area. a**, Regional growth rates in terms of surface area for the first (*top row*) and second (*bottom row*) postnatal years. **b**, ROI-specific mean laterality index for surface area.

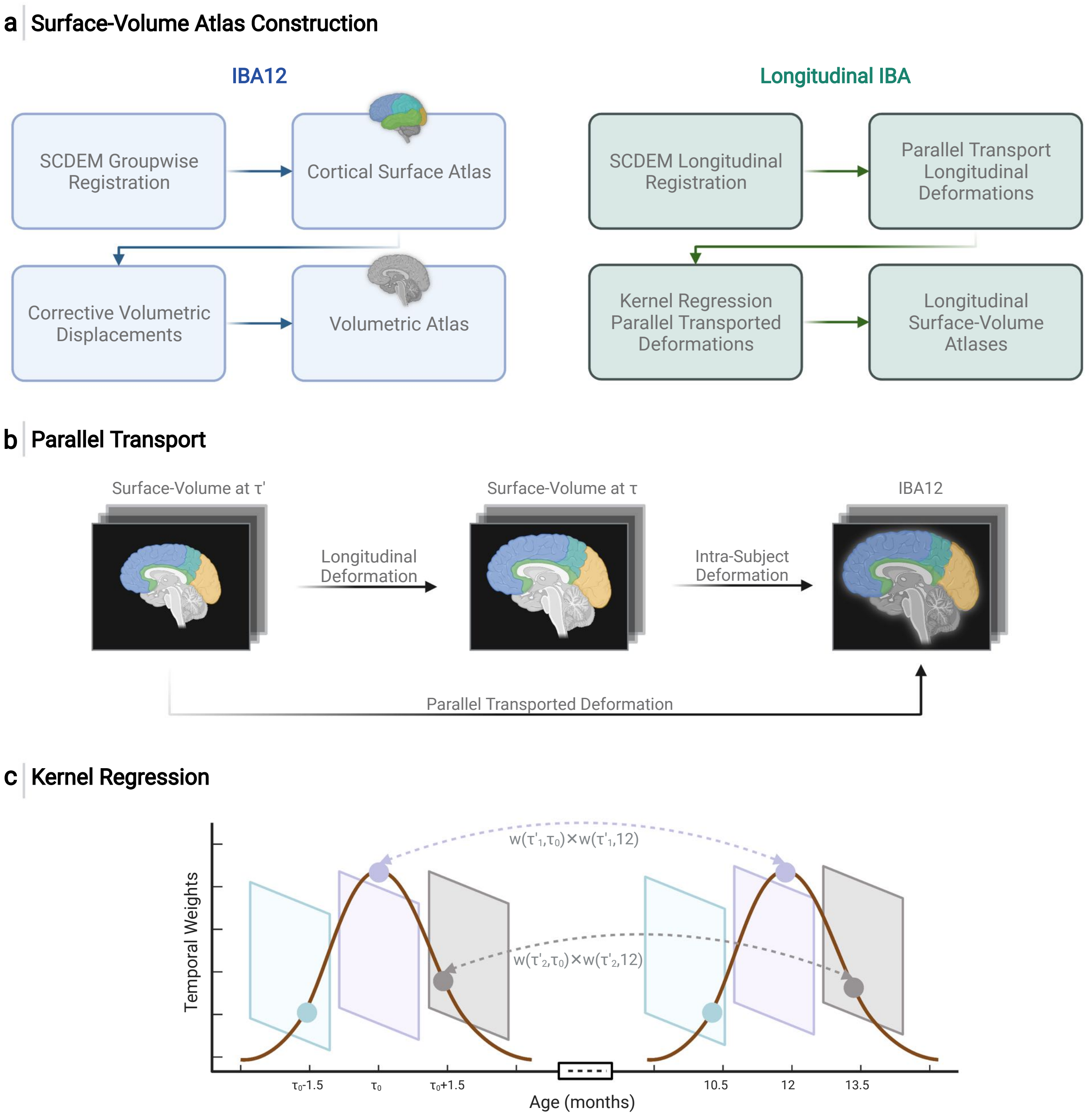


**Supplementary Fig. 20 | IBA construction pipeline.** **a**, Overview of the steps involved in constructing surface-volume IBA. **b**, Parallel transport of longitudinal deformations to the IBA12 space. **c**, Weighting of parallel transported deformations.

Supplementary Table 1. Statistical *t*-test of the laterality of average convexity.

| **Cortical ROI** | ***t*-score** | **DoF** |
|---|---|---|
| Caudal Middle Frontal | -42.68 | 24 |
| Entorhinal | -18.08 | 24 |
| Postcentral | -49.36 | 24 |
| Pars Triangularis | 7.84 | 20 |
| Supramarginal | -83.42 | 24 |
| Insula | -7.7 | 23 |
| Banks of Superior Temporal Sulcus | 32.71 | 24 |
| Lateral Orbitofrontal | 28.19 | 24 |
| Pars Orbitalis | 72.72 | 24 |
| Middle Temporal | -59.77 | 24 |
| Pericalcarine | -17.82 | 23 |
| Parahippocampal | -7.52 | 23 |
| Paracentral | 29.53 | 24 |
| Medial Orbitofrontal | 51.13 | 24 |
| Frontal Pole | 43.83 | 24 |
| Cuneus | 14.97 | 24 |
| Inferior Temporal | 8.88 | 21 |
| Rostral Middle Frontal | 39.03 | 24 |
| Rostral Anterior Cingulate | 14.95 | 24 |
| Isthmus Cingulate | 58.62 | 24 |
| Lateral Occipital | 104.51 | 24 |
| Lingual | -119.04 | 24 |
| Superior Parietal | -36.92 | 24 |
| Pars Opercularis | -12.44 | 24 |
| Fusiform | 6.76 | 21 |
| Caudal Anterior Cingulate | 80.51 | 24 |
| Superior Frontal | -68.45 | 24 |
| Temporal Pole | 4.63 | 13 |
| Precuneus | -59.82 | 24 |
| Transverse Temporal | 18 | 24 |
| Precentral | -13.84 | 24 |
| Inferior Parietal | 21.49 | 24 |
| Posterior Cingulate | 51.06 | 24 |
| Superior Temporal | -29.09 | 24 |

Supplementary Table 2. Statistical $t$-test of the laterality of mean curvature.

| Cortical ROI | $t$-score | DoF |
|---|---|---|
| Caudal Middle Frontal | 14.8 | 24 |
| Entorhinal | 21.4 | 24 |
| Postcentral | 78.4 | 24 |
| Pars Triangularis | 5.6 | 12 |
| Supramarginal | 103.2 | 24 |
| Insula | 26.3 | 24 |
| Banks of Superior Temporal Sulcus | 103.6 | 24 |
| Lateral Orbitofrontal | -18.8 | 24 |
| Pars Orbitalis | 7.7 | 17 |
| Middle Temporal | 83.7 | 24 |
| Pericalcarine | -27.7 | 24 |
| Parahippocampal | -9.7 | 24 |
| Paracentral | -25.6 | 24 |
| Medial Orbitofrontal | -23.9 | 24 |
| Frontal Pole | -19.2 | 24 |
| Cuneus | 49.0 | 24 |
| Inferior Temporal | 33.1 | 24 |
| Rostral Middle Frontal | 21.6 | 24 |
| Rostral Anterior Cingulate | 68.3 | 24 |
| Isthmus Cingulate | -37.7 | 24 |
| Lateral Occipital | -5.9 | 13 |
| Lingual | 56.2 | 24 |
| Superior Parietal | -17.5 | 24 |
| Pars Opercularis | 6.4 | 18 |
| Fusiform | -9.6 | 24 |
| Caudal Anterior Cingulate | -24.5 | 24 |
| Superior Frontal | -10.4 | 24 |
| Temporal Pole | 7.0 | 16 |
| Precuneus | 20.9 | 24 |
| Transverse Temporal | -20.8 | 24 |
| Precentral | 40.6 | 24 |
| Inferior Parietal | -24.1 | 24 |
| Posterior Cingulate | 18.9 | 24 |
| Superior Temporal | 68.6 | 24 |

Supplementary Table 3. Statistical *t*-test of the laterality of cortical thickness.

| Cortical ROI | *t*-score | DoF |
|---|---|---|
| Caudal Middle Frontal | -107.29 | 24 |
| Entorhinal | -36.92 | 24 |
| Postcentral | -61.36 | 24 |
| Pars Triangularis | 28.55 | 24 |
| Supramarginal | -139.52 | 24 |
| Insula | -21.83 | 24 |
| Banks of Superior Temporal Sulcus | -214.46 | 24 |
| Lateral Orbitofrontal | 29.84 | 24 |
| Pars Orbitalis | 37.23 | 24 |
| Middle Temporal | -88.45 | 24 |
| Pericalcarine | -236.35 | 24 |
| Parahippocampal | 29.13 | 24 |
| Paracentral | 20.14 | 24 |
| Medial Orbitofrontal | 47.57 | 24 |
| Frontal Pole | 44.24 | 24 |
| Cuneus | 19.24 | 24 |
| Inferior Temporal | -41.73 | 24 |
| Rostral Middle Frontal | 48.05 | 24 |
| Rostral Anterior Cingulate | 23.01 | 24 |
| Isthmus Cingulate | 85.07 | 24 |
| Lateral Occipital | 83.45 | 24 |
| Lingual | -106.03 | 24 |
| Superior Parietal | -102.53 | 24 |
| Pars Opercularis | -29.55 | 23 |
| Fusiform | -34.08 | 24 |
| Caudal Anterior Cingulate | 4.74 | 16 |
| Superior Frontal | -12.07 | 24 |
| Temporal Pole | -28.90 | 24 |
| Precuneus | -118.94 | 24 |
| Transverse Temporal | -12.21 | 23 |
| Precentral | -33.02 | 24 |
| Inferior Parietal | -57.37 | 24 |
| Posterior Cingulate | -39.48 | 24 |
| Superior Temporal | -37.44 | 24 |

Supplementary Table 4. Statistical $t$-test of the laterality of surface area.

| Cortical ROI | $t$-score | DoF |
|---|---|---|
| Caudal Middle Frontal | -6.6 | 24 |
| Entorhinal | -15.7 | 24 |
| Postcentral | -10.8 | 24 |
| Pars Triangularis | -37.5 | 12 |
| Supramarginal | -27.1 | 24 |
| Insula | -56.1 | 24 |
| Banks of Superior Temporal Sulcus | -28.7 | 24 |
| Lateral Orbitofrontal | 8.7 | 16 |
| Pars Orbitalis | -24.9 | 24 |
| Middle Temporal | -14.0 | 24 |
| Pericalcarine | -14.3 | 24 |
| Parahippocampal | -4.7 | 17 |
| Paracentral | -24.2 | 24 |
| Medial Orbitofrontal | 13.9 | 24 |
| Frontal Pole | -88.9 | 24 |
| Cuneus | -28.4 | 24 |
| Inferior Temporal | -5.0 | 14 |
| Rostral Middle Frontal | 13.1 | 23 |
| Rostral Anterior Cingulate | 129.6 | 24 |
| Isthmus Cingulate | 12.9 | 24 |
| Lateral Occipital | 13.6 | 24 |
| Lingual | -6.7 | 24 |
| Superior Parietal | -19.7 | 24 |
| Pars Opercularis | -4.2 | 12 |
| Fusiform | -4.9 | 20 |
| Caudal Anterior Cingulate | 36.5 | 24 |
| Superior Frontal | 11.1 | 19 |
| Temporal Pole | -24.2 | 24 |
| Precuneus | -30.5 | 24 |
| Transverse Temporal | 20.0 | 24 |
| Precentral | -13.4 | 24 |
| Inferior Parietal | -31.8 | 24 |
| Posterior Cingulate | 16.9 | 24 |
| Superior Temporal | -28.3 | 24 |

Supplementary Table 5. Neonatal and infant brain atlases currently available for neuroimaging studies.

| Study | Type | Components | Age Range | No. of Time Points |
|---|---|---|---|---|
| Kuklisova *et al.*[1] (2011) | Volumetric | T2w<br>Tissue map | 29 - 44 weeks GA[†] | 1 |
| Schuh *et al.*[2] (2018) | Volumetric | T1w & T2w<br>Tissue map | 35 - 44 weeks PMA[††] | 9 |
| Serag *et al.*[3] (2012) | Volumetric | T1w & T2w | 27 - 44 weeks PMA | 9 |
| Shi *et al.*[4] (2011) | Volumetric | T1w & T2w<br>Tissue map | 0 - 2 years | 3 |
| Sanchez *et al.*[5] (2012) | Volumetric | T1w & T2w | 2 weeks - 4 years | 13 |
| Zhang *et al.*[6] (2016) | Volumetric | T1w & T2w | 1 - 12 months | 5 |
| Oishi *et al.*[7] (2011) | Volumetric | T1w & T2w<br>DTI[†††] | 2 days | 1 |
| Kazemi *et al.*[8] (2007) | Volumetric | T1w | 39 - 42 weeks GA | 1 |
| Shi *et al.*[9] (2010) | Volumetric | T2w<br>Tissue map | 0.6 - 2 months | 1 |
| Hashioka *et al.*[10] (2011) | Volumetric | T2w | 1 week | 1 |
| Bozek *et al.*[11] (2018) | Surface | Fiducial cortical surfaces<br>Cortical features maps | 36 - 44 weeks PMA | 9 |
| Hill *et al.*[12] (2010) | Surface | Spherical cortical surfaces<br>Fiducial cortical surfaces<br>Sulcal depth map | $>$ 36 weeks GA | 1 |
| Wu *et al.*[13] (2019) | Surface | Spherical cortical surfaces<br>Cortical features maps<br>Cortical parcellation | 1 - 72 months | 11 |
| Li *et al.*[14] (2015) | Surface | Spherical cortical surfaces<br>Cortical features maps | 1 - 24 months | 7 |

[†] GA - gestational age [††] PMA - postmenstrual age [†††] DTI - diffusion tensor imaging
